# LRP2 contributes to planar cell polarity-dependent coordination of motile cilia function

**DOI:** 10.1101/2022.07.12.499714

**Authors:** Lena Bunatyan, Anca Margineanu, Camille Boutin, Mireille Montcouquiol, Sebastian Bachmann, Erik Ilsø Christensen, Thomas E. Willnow, Annabel Christ

## Abstract

Motile cilia are protruding organelles on specialized epithelia that beat in a synchronous fashion to propel extracellular fluids. Coordination and orientation of cilia beating on individual cells and across tissues is a complex process dependent on planar cell polarity (PCP) signaling. Asymmetric sorting of PCP pathway components, essential to establish planar polarity, involves trafficking along the endocytic path, but the underlying regulatory processes remain incompletely understood. Here, we identified the endocytic receptor LRP2 as regulator of PCP component trafficking in ependyma, a multi-ciliated cell type that is involved in facilitating flow of the cerebrospinal fluid in the brain ventricular system. Lack of receptor expression in gene-targeted mice results in a failure to sort PCP core proteins to the anterior or posterior cell side and, consequently, in the inability to coordinate cilia arrangement and to aligned beating (loss of rotational and translational polarity). LRP2 deficiency coincides with a failure to sort NHERF1, a cytoplasmic LRP2 adaptor to the anterior cell side. As NHERF1 is essential to translocate PCP core protein Vangl2 to the plasma membrane, these data suggest a molecular mechanism whereby LRP2 interacts with PCP components through NHERF1 to control their asymmetric sorting along the endocytic path. Taken together, our findings identified the endocytic receptor LRP2 as a novel regulator of endosomal trafficking of PCP proteins, ensuring their asymmetric partition and establishment of translational and rotational planar cell polarity in the ependyma.

## INTRODUCTION

The low-density lipoprotein receptor related protein 2 (LRP2) is a 600 kDa type-1 transmembrane protein of the LDL receptor gene family (Nykjaer and Willnow 2002). LRP2 is unique amongst this class of endocytic receptors as its expression in the mammalian organism is restricted to polarized epithelia, including the embryonic neuroepithelium as well as the renal proximal tubule, distal oviduct, epididymis, and ependyma in the adult (Christensen et al. 1995; Lundgren et al. 1997; Argraves and Morales 2004; Gajera et al. 2010; Willnow and Christ 2017). In these tissues, LRP2 localizes to the apical cell surface and subapical endocytic compartments, in line with its function in clearance of ligands by these absorptive epithelia. The physiological relevance of resorptive processes by this receptor is underscored by phenotypes seen in humans with familial *LRP2* deficiency (Donnai-Barrow/facio-oculo-acoustico-renal (DB/FOAR) syndrome and microforms of holoprosencephaly, HPE) featuring impaired development of the neuroepithelium into the forebrain (Kantarci et al. 2007; Rosenfeld et al. 2010). Comparable defects in forebrain development are seen in mouse models with induced *Lrp2* deficiency (Spoelgen et al. 2005).

So far, a role as endocytic receptor for the morphogen sonic hedgehog (SHH) is considered the main function of LRP2 during neuroepithelial development. By directing binding and cellular uptake of SHH by its receptor Patched 1, LRP2 facilitates SHH-dependent differentiation processes that govern transformation of the anterior neural tube into the forebrain (Christ et al. 2012; Christ et al. 2016). Interestingly, a recent study now suggests that the relevance of LRP2 for neuroepithelial development may not be restricted to endocytosis of SHH but include a role in neuroepithelial morphogenesis by directing dynamic remodeling of the apical cell compartment. Typically, alterations of this process lead to neural tube closure defects (NTDs), phenotypes shared by mouse (Kur et al. 2014; Sabatino et al. 2017) and frog models of LRP2 deficiency (Kowalczyk et al. 2021). NTDs in the latter models are linked to impaired apical cell constriction as well as loss of apical localization of Vangl2, a core planar cell polarity (PCP) protein (Kowalczyk et al. 2021).

In this study, we aim to provide further experimental support for a novel role of LRP2 in apical cell polarity by focusing on the action of this receptor in ependymal cells, a polarized epithelial cell type that lines the luminal surface of the brain ventricles (Redmond et al. 2019). With relevance to our hypothesis, ependymal cells are characterized by a bundle of motile cilia important for the generation of cerebrospinal fluid (CSF) flow (Worthington and Cathcart 1963; Kumar et al. 2021). These cilia consist of a basal segment anchored to the apical cell membrane and an axoneme, extending from the cell surface into the ventricular lumen. The basal segment of the cilium is composed of the basal body (BB), associated with the basal foot, transitional fibers, and striated rootlets. Besides a role in cell surface anchoring, the basal apparatus determines the beating direction of the cilia. This direction matches the effective stroke direction of the ciliary shaft, providing coordinated beating of a ciliary bundle on a cell (Satir et al. 2014).

Coordination of motile cilia localization and beating on ependyma cells is a process intimately linked to the PCP pathway. Alignment of BB orientation in ependymal cells is initiated in their precursor cells, the radial glia around embryonic (E) day 16.5. Postmitotic radial glial cells are characterized by a single primary cilium. Around E16.5, the primary cilium becomes asymmetrically displaced on the apical surface and translational polarity is established (Mirzadeh et al. 2010). This initial displacement is driven by a passive weak CSF flow generated by fluid secretion in the choroid plexus and absorption in the foramen of Monroe, defining CSF flow in a caudo-rostral direction. Around P2, immature ependymal cells have grown a bundle of initially widely scattered and randomly beating cilia (Boutin et al. 2014; Ohata and Alvarez-Buylla 2016). At this time point, the asymmetric localization of core PCP proteins into two clusters on opposite sides in the apical plasma membrane compartment becomes apparent (Yang and Mlodzik 2015; Butler and Wallingford 2017). For example, Van Gogh-like 2 (Vangl2) and cadherin epidermal growth factor (EGF) like laminin G-like seven-pass G-type receptor 1 (Celsr1) localize to the posterior cell membrane, opposite to the side of ciliary patch displacement. Cytosolic adaptor proteins like Daple (Ccdc88c) and Dishevelled1 (Dvl1) localize to the anterior cell membrane, the side the ciliary patch is displaced to (Harrison et al. 2020). This asymmetric localization of PCP proteins is a prerequisite for BBs to align in a common orientation and to beat synchronously in individual cells as well as across the entire ependyma (tissue-level planar polarity). As a result, rotational polarity is established, generating an active CSF flow (Guirao et al. 2010; Wallingford 2010; Ohata and Alvarez-Buylla 2016). As PCP defects commonly result in motile cilia dysfunction (Kumar et al. 2021; Hyland and Brody 2021; Kapania et al. 2022), we performed structural and functional characterization of motile cilia in the LRP2-deficient ependyma to gain further insights into a possible role for LRP2 in PCP pathway function, and the underlying molecular mechanisms of receptor action.

## MATERIALS AND METHODS

### Mouse models

LRP2 deficient *Lrp2^-/267^* mice were generated by crossing heterozygous mice with targeted *Lrp2* gene defect (Willnow et al. 1996) with heterozygous mice identified from an ENU screen for mutations impairing brain cortex morphogenesis (Zarbalis et al. 2004) carrying a T to A transition at amino acid position 2721 resulting in a stop codon. The combination of both null allele results in a complete *Lrp2* gene defect analyzed in *Lrp2^-/267^*mice in comparison to either wild-type (*Lrp2^+/+^*) or heterozygous for one of the mutant *Lrp2* allele (*Lrp2^+/-^; Lrp2^+/267^*). No LRP2 haploinsufficiency phenotype was detected in *Lrp2^+/-^* and *Lrp2^+/267^* mice. Together with *Lrp2^+/+^* mice these animals were combined and referred to as control group in this study. Adult (>P70) and juvenile (P20-P40) mice were used in experiments as indicated.

### Immunhistochemical analysis

Whole mount dissection of the ventricular lateral wall (LW) and following immunostainings were performed as described previously (Mirzadeh et al. 2008; Mirzadeh et al. 2010). In brief, freshly dissected whole mount LW preparations were washed for 1 min in 0.1% Triton X-100 in 1x DPBS, afterwards fixed for 10 min in methanol at −20°C, followed by washing 3 times in 0.1% Triton X-100 in 1x DPBS and incubated for 1 hr in PBS and 3% BSA blocking solution at RT. Primary antibodies were diluted in blocking solution and whole mount LW preparations were incubated over night at 4°C. The following primary antibodies were used: mouse IgG2b anti-acetylated-a-tubulin (1:1000; Sigma), mouse IgG1 anti-g-tubulin(1:400; Abcam), mouse IgG1 anti-b-catenin (1:500; BD Biosciences), mouse IgG2b anti-FGFR1OP (FOP; 1:1500; Abnova), rabbit anti-ZO1 (1:500; Invitrogen), rabbit anti-Arl13b (1:250; Proteintech), rabbit anti-Daple (1:100; IBL), rabbit anti-NHERF1 (1:300; Alomone labs), rabbit anti-CASMAP2 (1:100; Proteintech), rat anti-EB3 (1:100; Abcam), mouse anti-Dvl1 (1:100; Santa Cruz Biotechnology), rabbit anti-Vangl2 (1:300; gift from Mireille Montcouquiol, Neurocenter Magendie, Bordeaux, France), rabbit anti-Celsr1 (1:1000; gift from Elaine Fuchs, The Rockefeller University, New York, USA), guinea pig anti-LRP2 (1:1000; homemade). Following washing in 0.1M DPBS, primary antibodies were visualized using secondary antisera conjugated with Alexa Fluor 405, 488, 568, and 647 (1:500; Invitrogen and Jackson Immuno Research), or with Biotin-SP (1:100; Jackson Immuno Research) followed by fluorescent conjugates of streptavidin 647 (1:500; Invitrogen) in 0.1M DPBS. Finally, whole mount LW preparations were mounted using Dako fluorescent mounting medium. Image acquisitions were carried out with a Leica TCS SP8 confocal microscope using a 63x PL APO CS2 oil immersion objective (NA 1.4). The excitation lines corresponding to the Alexa dyes used here were provided by a white light laser and a 405 nm diode laser and were separated from the fluorescence emission using an acousto-optic beam splitter (AOBS). The fluorophores emission was collected in specific bands via a spectral detector and directed on 3 photomultiplier tubes. Sequential excitation/emission scanning was used to avoid cross-talk between fluorophores. Images were acquired using a pixel size of 90 nm and the z planes were separated by 1 µm.

### Data quantification of ciliary patch organization

To quantify cilia beating parameters whole mount LW preparations were immunostained for g-tubulin,FOP and ZO1 as described above and the Biotool1 software developed by Paul Labedan and Camille Boutin (Labedan et al. 2016) was used for further analyses as described previously (Boutin et al. 2014). To investigate patch displacement, the software was used to manually trace the contours of individual ependymal cells and their respective ciliary patches or primary cilia patches. The geometric centers of the marked areas were calculated and two vectors for each cell, one describing the basal body (BB) patch displacement relative to the center of each cell, and one describing the mean vector of displacement in the entire microscopic field were defined in the software. The angle resulting from those two vectors of each cell was defined as VpatchD reflecting the patch displacement in the investigated field. To analyze beating orientation of a motile cilia patch the vector connecting FOP to g-tubulin for individual cilia of a patch was manually drawn and the mean vector of all vectors in the analyzed field is calculated in the software. The angle between those two vectors is defined as VpatchO (mean vCil) and describes the coordination of cilia orientation between all cells in the investigated field. To determine the coordination of patch displacement with patch beating orientation from the cell center, the angles between VPatchD and VPatchO vectors (VpatchD&O) were quantified. In addition, individual cilium beating coordination within a cell was identified by vCil vectors and used to calculate the circular standard deviation (CSD) value for each cell. A higher CSD value indicates more non-coordinated beating of individual cilia within a cell. The extent of the patch displacement relative to center of the cell (strength) was determined by an algorithm in Biotool1 software. Furthermore, Biotool1 software counted the cilia numbers in each cell.

### Electron microscopy

For immunoelectron microscopy, whole mount LW tissue was immersion-fixed in 3% paraformaldehyde/0.05% glutaraldehyde buffered in PBS, freeze-substituted, and embedded in London LR White hydrophilic resin. Ultrathin sections were cut, placed on Ni-grids, incubated with rabbit anti-LRP2 (1:5,000; homemade) antisera followed by immunogold particle-coupled secondary antibody (Dako Glostrup), and studied in a Zeiss EM906 transmission electron microscope. For scanning electron microscopy, whole mount LW tissue was fixed in 2.5% PBS-buffered glutaraldehyde and dehydrated in alcohol, osmicated, dried in a critical-point apparatus and coated with carbon for examination on a Zeiss scanning electron microscope. For transmission electron microscopy, tissues were fixed in 1% glutaraldehyde in PBS, following by postfixation in 1% OsO4 (in sodium cacodylate buffer). After stained en bloc in saturated uranyl acetate, the tissue was dehydrated in a graded ethanol series and embedded in Epon. Sections were cut with a Leica Ultramicrotome UCT, stained with uranyl acetate and lead citrate for examination in a FEI 100 CM electron microscope.

### Ciliary beat frequency analysis

Freshly dissected LW whole mount preparations from 6 wild-type and 4 *Lrp2^-/267^* mice were incubated with rat anti-CD24 conjugated with Phycoerythrin (PE; 1:100; BD Pharmingen) in Neurobasal medium (Gibco) supplemented with B-27 serum-free supplement (Gibco) for 20 min at RT. After washing with L-15 medium (Gibco) whole mount LW preparations were placed in glass-bottomed dishes (ibidi) within the cavity delimited by two spacers glued on top of each other (0.12 mm deep; Invitrogen) and covered by a cover slip (Thermo Scientific), with the tissue surface facing the glass bottom. Ciliary beating was imaged with an inverted wide field microscope (Olympus) using a 60x water immersion lens (NA 1.1, working distance 1.5 mm) at 37°C, under 5% CO_2_. The emission of a Lumencor Spectra X source was selected with a 575/25 nm bandpass filter to excite the PE, while fluorescence was collected via a 623/24 nm filter. Images were acquired as time lapses of 600 frames using an EMCCD camera (Hamamatsu) with an integration time per frame of 10 ms, leading to an imaging speed of 21 frames/s when considering also the readout time of the whole camera chip (512×512 pixels). Depending on the size of the dissected tissue fragments, 5-20 different fields of view were imaged for each sample. Raw time lapse images were processed using background subtraction, bleach correction and smoothing functions available in ImageJ/Fĳi (Miura 2020). On each time lapse, 5-12 cilia were selected using a line, and a kymogram was constructed via the Multi Kymograph routine available in ImageJ/Fĳi. The fluorescence intensity oscillation along a kymogram image was exported as intensity vs. time graph, which was further processed in Python using a custom written routine to calculate the fast Fourier transform (FFT). The frequency corresponding to the dominant peak determined via thresholding of the FFT spectrum was considered as the beating frequency.

### Statistical analysis

For classical statistics, two-tailed Student’s t test was applied using Graph Pad Prism 7. All data were presented as standard error of mean (SEM). For circular statistics, controls and LRP2-deficient genotypes were compared applying Watson’s U2 test using the circular statistics software program Oriana. Difference in data were considered significant with p<0.05.

## RESULTS

### LRP2 localizes to the ciliary pocket and basal body of motile cilia

To interrogate a possible role for LRP2 in structural and functional integrity of motile cilia, we made use of a mouse model compound heterozygous for two mutant *Lrp2* allele, a receptor null allele (*Lrp2^-^*) generated by homologous recombination (Willnow et al. 1996) and a missense mutation obtained in an ENU screen (*Lrp2^267^*) (Zarbalis et al. 2004). Both alleles cause complete loss of LRP2 activity and show identical receptor deficiency phenotypes. For reasons attributed to distinct genetic modifiers in both lines, animals compound heterozygous for the two mutant alleles (*Lrp2^-/267^)* show improved perinatal survival as compared to the respective homozygous mutant lines, enabling us to generate sufficient adult mutant mice for analyses. No phenotypic differences are seen comparing wild-type (*Lrp2^+/+^*) and heterozygous animals (*Lrp2^+/-^* or *Lrp2^+/267^*) (Gajera et al. 2010). Therefore, both genotypes were used as reference for receptor null animals (*Lrp2^-/267^),* collectively referred to as control animals in this study.

Expression of LRP2 on the apical surface of ependymal cells has been described by us before (Gajera et al. 2010). Here, we refined the expression analyses by focusing on the subcellular localization of the receptor with respect to the motile cilia in this cell type. To do so, we stained *en face* preparations of the lateral wall of murine wild-type ependyma for LRP2 as well as for γ-tubulin and acetylated-tubulin, markers of basal foot processes and shaft of motile cilia, respectively (Hagiwara et al. 2000). LRP2 immunosignals localized to the ciliary patch region on the apical cell surface, in close proximity to the basal foot marker γ-tubulin (Fig. 1A). Localization of LRP2 close to the ciliary patch was substantiated by immunoelectron microscopy, documenting its presence at the apical cell surface and in subapical endosomes of the ciliary pocket, an area highly active in endocytosis (Molla-Herman et al. 2010) (Fig. 1B, arrowheads). Loss of LRP2 did not impact the structural appearance of motile cilia as documented by scanning and transmission electron microscopy (Fig. 1C). In addition, immunostaining of basal foot marker γ-tubulin, BB marker FOP (FGFR1 oncogene partner) as well as ciliary shaft markers Arl13b and acetylated tubulin confirmed comparable cilia structure in *Lrp2^-/267^* and control mice (Fig. 1D).

**Figure 1.**
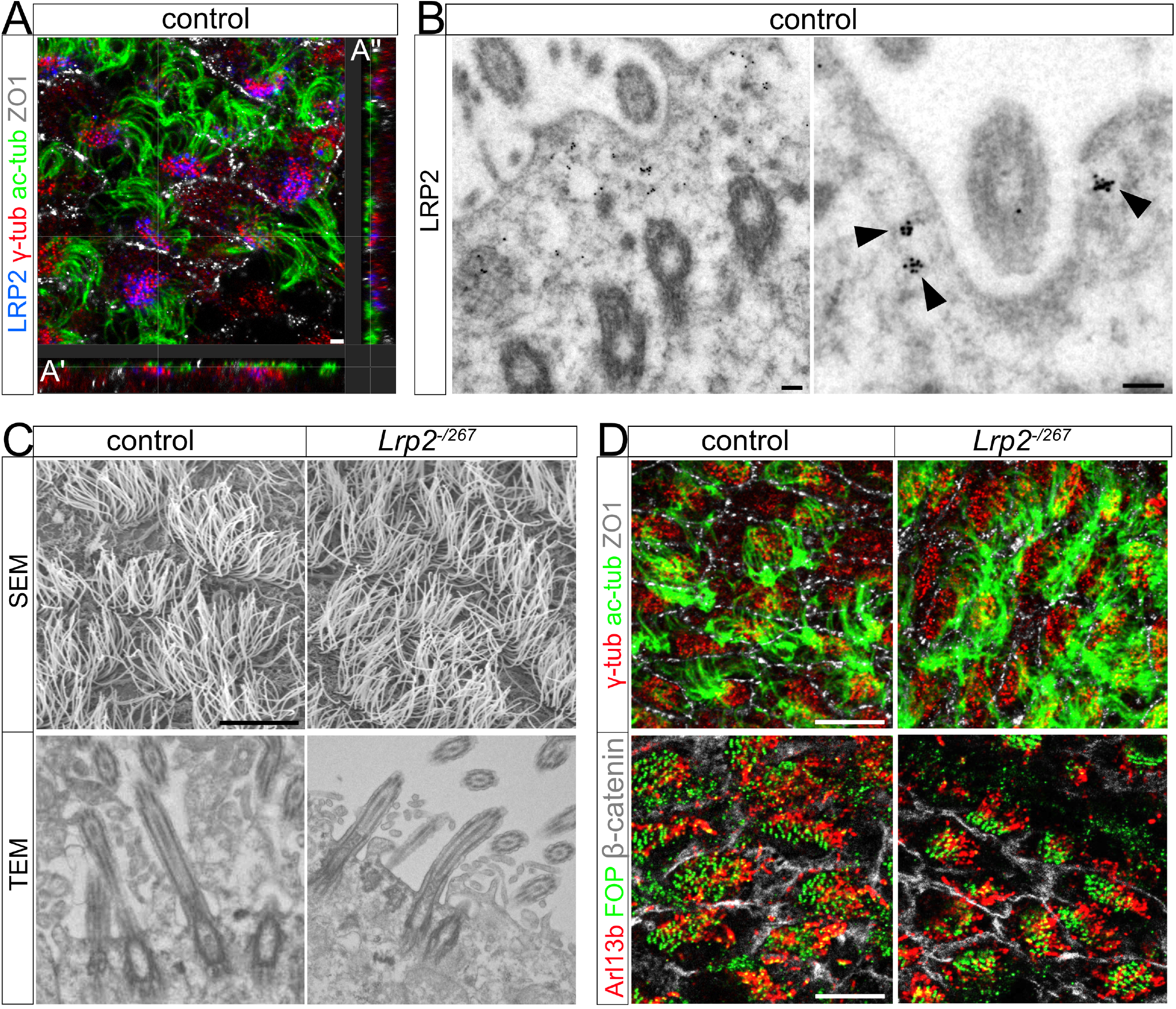
LRP2 localizes to the ciliary patch region of ependymal cells. **(A)** Immunohistological detection of LRP2 (blue), basal body marker γ-tubulin (red), ciliary shaft marker acetylated-tubulin (green), as well apical cell surface marker zonula occludens-1 (ZO1, grey) on *en face* preparations of the ventricular lateral wall of adult wild-type mice. LRP2 abundantly localizes to the ciliary patch region of motile cilia. Insets A’ and A’’ depict images in z-axis configuration highlighting close proximity of LRP2 to g-tubulinat the basal foot below the ciliary shaft (marked by acetylated-tubulin). Scale bar: 2 µm. (**B**) Immunoelectron microscopical detection of LRP2 on the apical cell surface of the adult ependyma. The high magnification inset documents LRP2 localizing to the apical cell membrane and subapical endosomes of ciliary pocket (arrowheads). Scale bar: 100 nm. (**C**) Scanning electron microscopic (SEM) images of the apical ependymal surface in control and *Lrp2^-/267^* mice documenting comparable appearance of motile cilia bundles in the ependyma of *Lrp2^-/267^*and control mice. Scale bar: 10 µm. Also, transmission electron microscopy (TEM) pictures show identical molecular architecture of motile cilia in both genotypes. (**D**) Immunohistological analyses of *en face* lateral wall preparations stained for g-tubulin(red), acetylated-tubulin (green) and ZO1 (grey) in the upper panel and for Arl13b (red), FGFR1 oncogene partner (FOP, green) and b-catenin (grey) in the lower panel. No structural differences could be detected between the two genotypes. Scale bar: 10 µm.

### LRP2 deficiency impacts ciliary patch displacement in the murine ependyma

To evaluate a potential impact of LRP2 deficiency on the functional integrity of motile cilia we employed an experimental strategy to co-immunostain *en face* preparations of the lateral ventricular wall for ciliary BB marker FOP, ciliary basal foot marker γ-tubulin, as well as the apical cell surface marker Zonula Occludens-1 (ZO1) (Boutin et al. 2014; Labedan et al. 2016), (Fig. 2A, upper panel). Using the Biotool software (github.com/pol51/biotool1), the contours of individual ependymal cells (based on ZO1 signals) and their respective ciliary patches (based on FOP signals) were traced and used to calculate various morphological parameters, including the ciliary patch size and its displacement relative to the center of the cell (i.e., translational polarity). As the basal foot protruding from the basal body points in the direction of the cilia stroke (Marshall and Kintner 2008), the localization of immunosignals for FOP and γ-tubulin relative to each other could also be used to accurately document the beating direction of individual cilia by drawing vectors from FOP to γ-tubulin immunosignals for each cilium (Vcil), (Fig. 2A, lower panels). These vectors indicate the directionality of beating of individual cilia within a patch (CSD value) and also the directionality of beating of patches on individual cells or across the entire field (rotational polarity).

**Figure 2.**
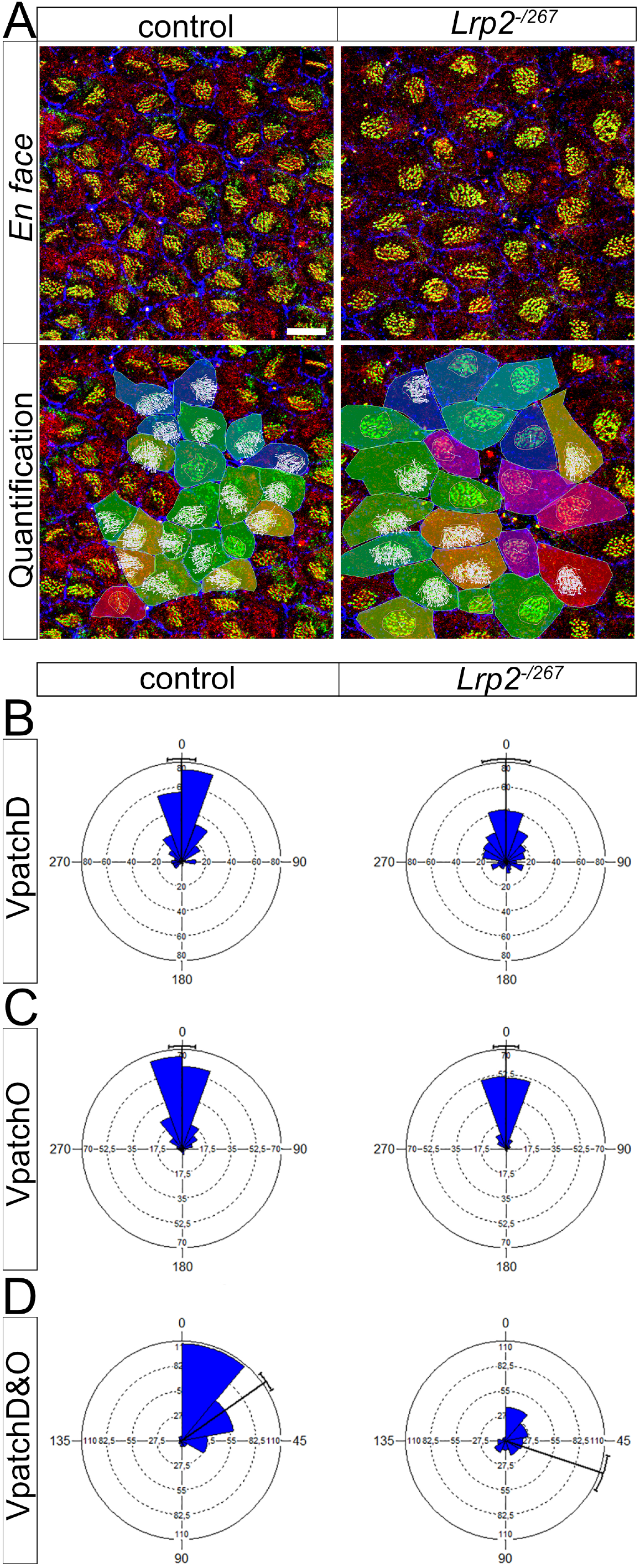
Misalignment of ciliary patch displacement and beating orientation in the LRP2-deficient juvenile ependyma. **(A)** The upper panels show *en face* views of the ventricular lateral wall of juvenile control and *Lrp2^-/267^* mice (postnatal day 20-28) immuno-stained for basal body markers FOP (green) and γ-tubulin (red), as well as for apical cell surface marker ZO1 (blue). Scale bar: 10 µm. The lower panels depict the same images visualizing analysis of tissue-wide planar cell polarity using the Biotool1 software (github.com/pol51/biotool1). White arrows indicate the vectors from FOP to γ-tubulin immunosignals in individual cilia, defined as Vcil. Color coding of individual cells describes the degree of ciliary patch displacement relative to the centre of the cell. The stronger the deviation of patch displacement of individual cells from the average vector of the entire field, the further the colour shifts from green towards the red or blue colour spectra. In cells without arrows, VpatchO could not be defined because the maximum circular standard deviation (CSD) value was above the threshold set at 45 (see methods for details). (**B-D**) Graphical representation of circular statistical analysis using Watson U^2^ test (controls: 265 cells, *Lrp2^-/267^* mutants: 284 cells, 5 animals per genotype) documenting impaired coordination of ciliary patch displacement (VpatchD, p<0.001; B) as well as beating orientation (VpatchO, p<0.01; C) in receptor-deficient as compared to control cells. Also, alignment of patch displacement and beating orientation (VpatchD&O; D) is lost in mutants as shown by Watson U^2^ test (controls: 220 cells, *Lrp2^-/267^*: 139 cells, 5 animals per genotype; p<0.001).

Initially, we evaluated the translational polarity in control and mutant ependyma of juvenile mice by determining the coordinated displacement of ciliary patches from the center of individual cells. As depicted in immunostained *en face* preparations in Fig. 2A, ependymal cells in the control tissue manifested a tissue-wide coordinated and directed displacement of the ciliary patches relative to cell center. By contrast, many ependymal cells in the LRP2-deficient tissue showed random displacement of their ciliary patches. Impaired ciliary patch displacement in LRP2-deficient ependyma was substantiated by calculating the integrated angle values VpatchD (Fig. 2B). They describe the mean distribution of ciliary patch displacement for each cell relative to the mean vector of displacement in the entire field of observation. In controls, most of vectors were distributed between −45° and 45° around the mean, demonstrating coordination of ciliary patch displacement. In *Lrp2^-/267^*ependyma, coordination of patch displacement was partially lost, as documented by a broader distribution of vectors from individual cells within the field. Circular statistical analysis by Watson U^2^ test confirmed significant differences in ciliary patch displacement when comparing both genotype groups (p<0.001; Fig. 2B).

A prerequisite for correct ciliary patch displacement in mature ependymal cells and establishment of proper translational polarity is the displacement of the primary cilium in progenitors of ependymal cells, the radial glia cells (Mirzadeh et al. 2010). We detected coordinated displacement of the primary cilium in immature ependymal cells of control neonates using immunostaining for γ-tubulin and ZO1 (Fig. S1A) and circular statistical analysis of the corresponding data (Fig. S1B). By contrast, VpatchD angles in *Lrp2^-/267^* newborns showed significantly broader distribution around the mean (Fig. S1B, p<0.001).

To test how LRP2 activity impacts the tissue-level planar polarity and coordination of cilia beating orientation established in mature ependymal cells, we used the Vcil data of individual cilia from juvenile animals (Fig. 2A) to determine the VpatchO angle values. They describe the beating orientation of the motile cilia in each cell with respect to the entire field of analysis. Circular distribution of VpatchO around the mean displayed a broader distribution in juvenile *Lrp2^-/267^* mice compared to controls, demonstrating a statistically significant decrease in coordination of ciliary beating orientation in mutant ependyma (p<0.01; Fig. 2C). According to earlier reports, beating orientation and patch displacement direction are aligned in ependymal cells of the adult murine brain (VpatchD&VpatchO vectors; Boutin et al., 2014). In line with defects in both patch displacement and beating orientation, VpatchD&VpatchO vectors indicated random directions in juvenile LRP2 mutants compared with controls, as reflected in a wider range of angle values around the mean in the circular dispersion graphs (p<0.001; Fig. 2D). Defects in ciliary displacement and beating orientation were also confirmed in brains of adult mutant mice (P >70) by determination of vectors for VpatchD, VpatchO, and VpatchD&VpatchO (Fig. S2).

### LRP2 deficiency impacts rotational polarity but not ciliary beat frequency of the murine ependyma

Next, we investigated the impact of LRP2 deficiency on coordination of cilia beating within a bundle (i.e., rotational polarity). To do so, we focused on the analysis of ependymal cells during juvenile stages (P20-35) when ependymal cells acquired their proper planar cell polarity (PCP), and demonstrated correct positioning of PCP proteins (Guirao et al. 2010; Ohata and Alvarez-Buylla 2016). Rotational polarity in control tissue was confirmed by low circular standard deviation (CSD) values for vectors Vcil (Fig. 3A and B). By contrast, CSD values where significantly higher in *Lrp2^-/267^* ependymal cells, documenting impaired rotational polarity in cells lacking LRP2 (Fig. 3B, p<0.0001). This defect was also seen when displaying CSD values in frequency intervals of 10, showing a shift from intervals with lower CSD values in controls to intervals with higher CSD values in mutant cells (Fig. 3C, p<0.0001). Similar defects in rotational polarity were confirmed in the ependyma of adult *Lrp2^-/267^* mice (Fig. S3). Changes in translational and rotational polarity in *Lrp2^-/267^* ependyma were likely not due to morphological abnormalities in cilia patches as cilia number per patch (Fig. S4A), apical cell surface area (Fig. S4B), patch area (Fig. S4C), or the extent of patch displacement from the center (strength, Fig. S4D) were comparable between control and receptor-deficient juvenile ependyma.

**Figure 3.**
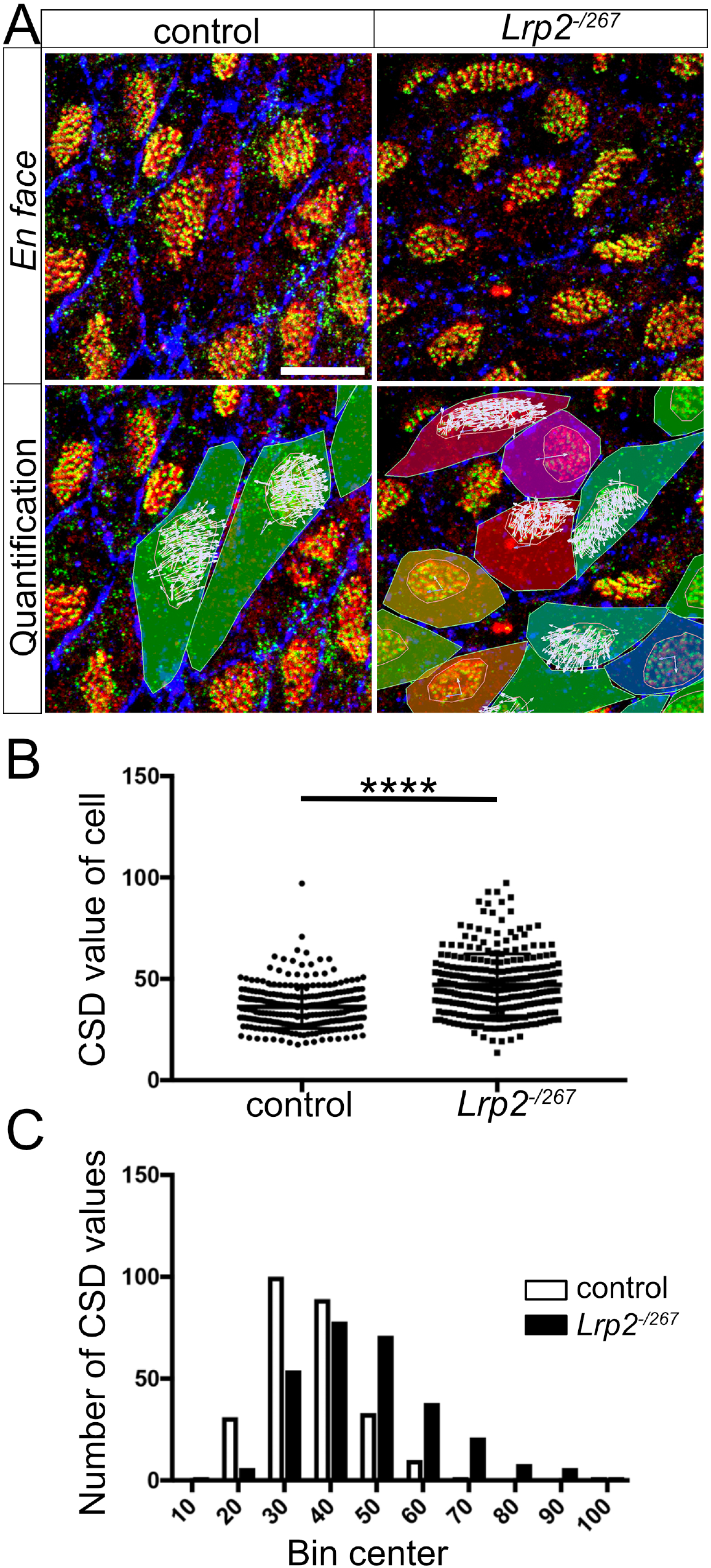
Defects in rotational polarity in LRP2-deficient juvenile ependymal cells. (**A**) The upper panels show *en face* preparations of the ventricular lateral wall of juvenile control and *Lrp2^-/267^*mice (postnatal day 20-28) immuno-stained for FOP (green), γ-tubulin (red), and ZO1 (blue). Scale bar: 10 µm. The lower panels show the same images visualizing the vectors Vcil (white) from FOP to γ-tubulin immunosignals, indicative of the beating orientation of individual cilia. Color coding of individual cells describes the degree of ciliary patch displacement relative to the centre of the cell (with stronger deviation of patch displacement of individual cells from the average vector of the entire field marked in red or blue). (**B**) Circular standard deviation (CSD) values for individual cilia in control and *Lrp2^-/267^* juvenile ependyma (control: 265 cells, *Lrp2^-/267^*: 284 cells; 5 animals per genotype). CSD values are significantly higher in *Lrp2^-/267^*mice as determined by unpaired *t*-test (p<0.0001). (**C**) CSD values for data in panel B grouped into frequency intervals of 10 (from 0 – 100), demonstrating enrichment of intervals with lower CSD values in control cells as compared with intervals with higher CSD values in *Lrp2^-/267^* cells.

Defects in rotational polarity by disturbed coordinated beating of motile cilia can affect ciliary beating frequency (CBF) (Park et al. 2008). Therefore, we investigated CBFs in control and *Lrp2^-/267^* adult ependyma by labelling cilia with the phycoerythrin (PE)- conjugated glycosyl phosphatidylinositol-anchored sialoglycoprotein CD24. Video recordings of beating cilia on *en face* preparations of both genotypes were used as basis for quantifying ciliary beat frequency in cycles per second (Fig. 4A, companion files: video control and video LRP2 mutant). In detail, motile cilia were marked with a line to determine the corresponding kymogram of the moving cilium (Fig. 4A, orange line). No significant differences were observed between control and *Lrp2^-/267^*mice in the kymogram patterns visualizing the time variation of fluorescence intensity (Fig. 4 compare B^1^ with C^1^). Following, the kymograms were converted into graphs visualizing the fluorescence intensity over time (Fig. 4B^2^, C^2^). Using fast Fourier transform (FFT), the mean beating frequency of individual cilia was determined (Fig. 4B^4^, C^4^). CBFs in control (mean: 4.96; SD: 0.4) and *Lrp2^-/267^*(mean: 5.17; SD: 0.4) ependyma were not significantly different (Fig. 4D). Thus, while beating of cilia is uncoordinated in LRP2-deficient ependyma, the speed of ciliary beating is normal.

**Figure 4.**
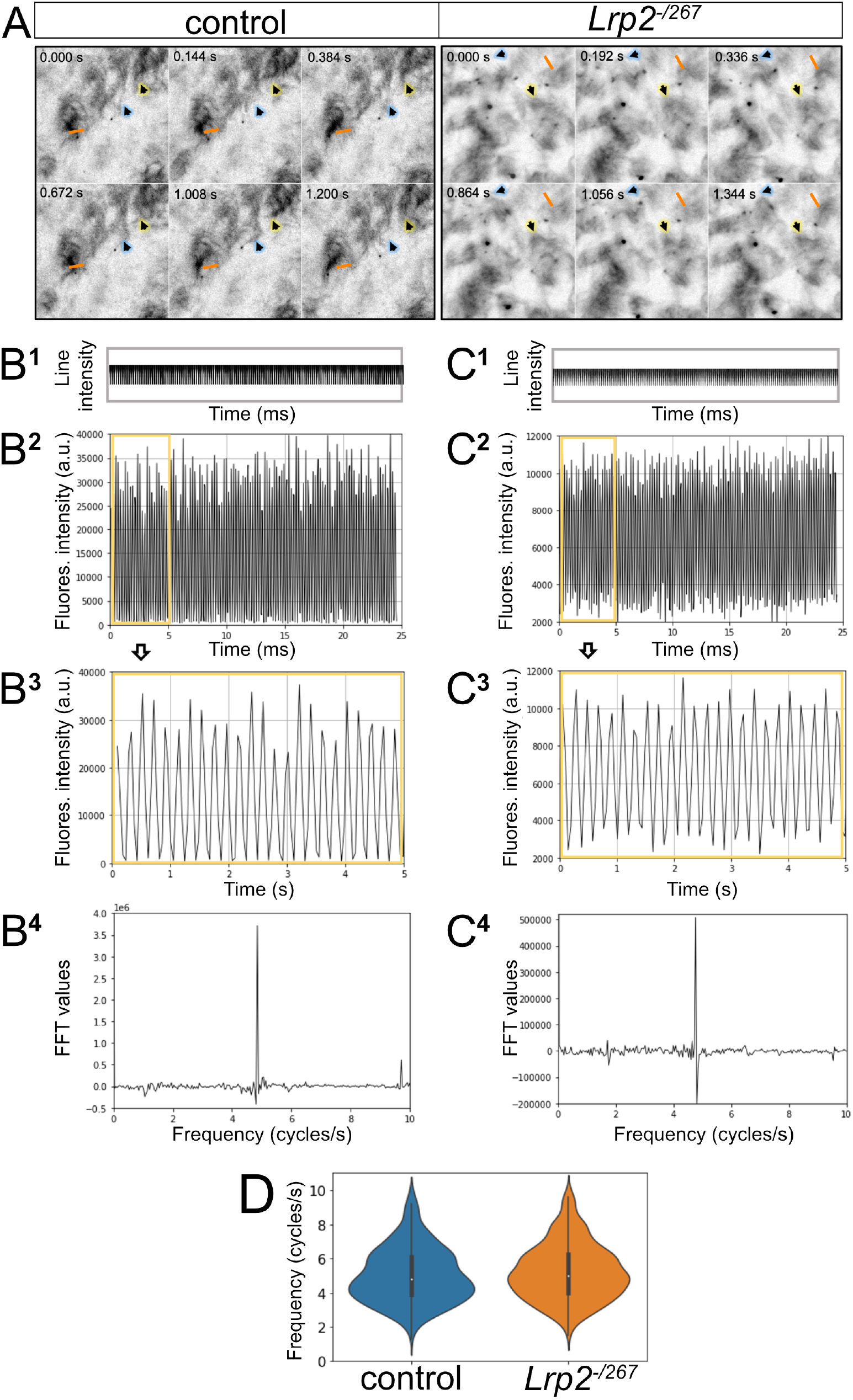
Beating frequency of motile cilia in control and LRP2 mutant ependyma. (**A**) Images of cilia movement were acquired as time lapses and examples are shown at the indicated time points. Regions of interest (ROIs) were marked as lines on patches of beating cilia (orange lines). Arrows indicate other examples of patches of moving cilia during image acquisition. Scale bar: 10 µm. (**B^1^**) Typical kymogram obtained from a control mouse by selecting a moving cilium or group of cilia, showing the line intensity on each consecutive image in the time series. (**B^2^**) Time variation of the fluorescence intensity along the kymogram displayed as graph. (**B^3^**) The evolution of the graph between 0-5 s is enlarged for better visualization. (**B^4^**) The fast Fourier transform (FFT) spectrum obtained from the depicted time trace in B^3^. The dominant peak indicates the beating frequency of the selected cilium. (**C^1-4^**) The same features as in B shown for a cilium from a *Lrp2^-/267^* mouse sample to illustrate that no significant differences were observed when compared to the control. (**D**) Violin plots demonstrate no significant change in beating frequency of motile cilia comparing control (408 cells) and *Lrp2^-/267^* (735 cells) mice.

### Loss of LRP2 impairs proper localization of core PCP proteins in ependymal cells

Previous studies demonstrated the importance of cytoskeletal rearrangements in establishing the cilia polarity (Hirota et al. 2010; Werner et al. 2011; Vladar et al. 2012; Butler and Wallingford 2017). In mature ependymal cells, microtubules underlie the BB and extend between the ciliary patch and the cell cortex, anchoring cilia to the cytoskeleton (Hirota et al. 2010; Kunimoto et al. 2012; Boutin et al. 2014).

To evaluate possible defects in microtubule architecture in LRP2-deficient ependyma, we immunostained *en face* preparations of the lateral ventricular wall of control and of *Lrp2^-/267^* juvenile mice for markers of plus- and minus ends of microtubules (MT). The microtubule plus-end-tracking protein end-binding protein 3 (EB3) is a molecular link between MTs and the actin cytoskeleton (Geraldo et al. 2008; Jaworski et al. 2009). It localized in a coordinated fashion to the anterior side of ependymal cells in control tissue (Fig. 5A, arrows). A similar EB3 localization was seen in *Lrp2^-/267^* mice ependyma. Immunostaining for calmodulin-regulated spectrin-associated protein 2 (CAMSAP2), a MT minus-end targeting protein (Akhmanova and Hoogenraad 2015; Robinson et al. 2020; Liu et al. 2021), also showed comparable subcellular localization in control and *Lrp2^-/267^* tissue (Fig. 5B). Jointly, these findings argued against defects in the MT architecture as the molecular cause of impaired polarity in the LRP2-deficient ependyma.

**Figure 5.**
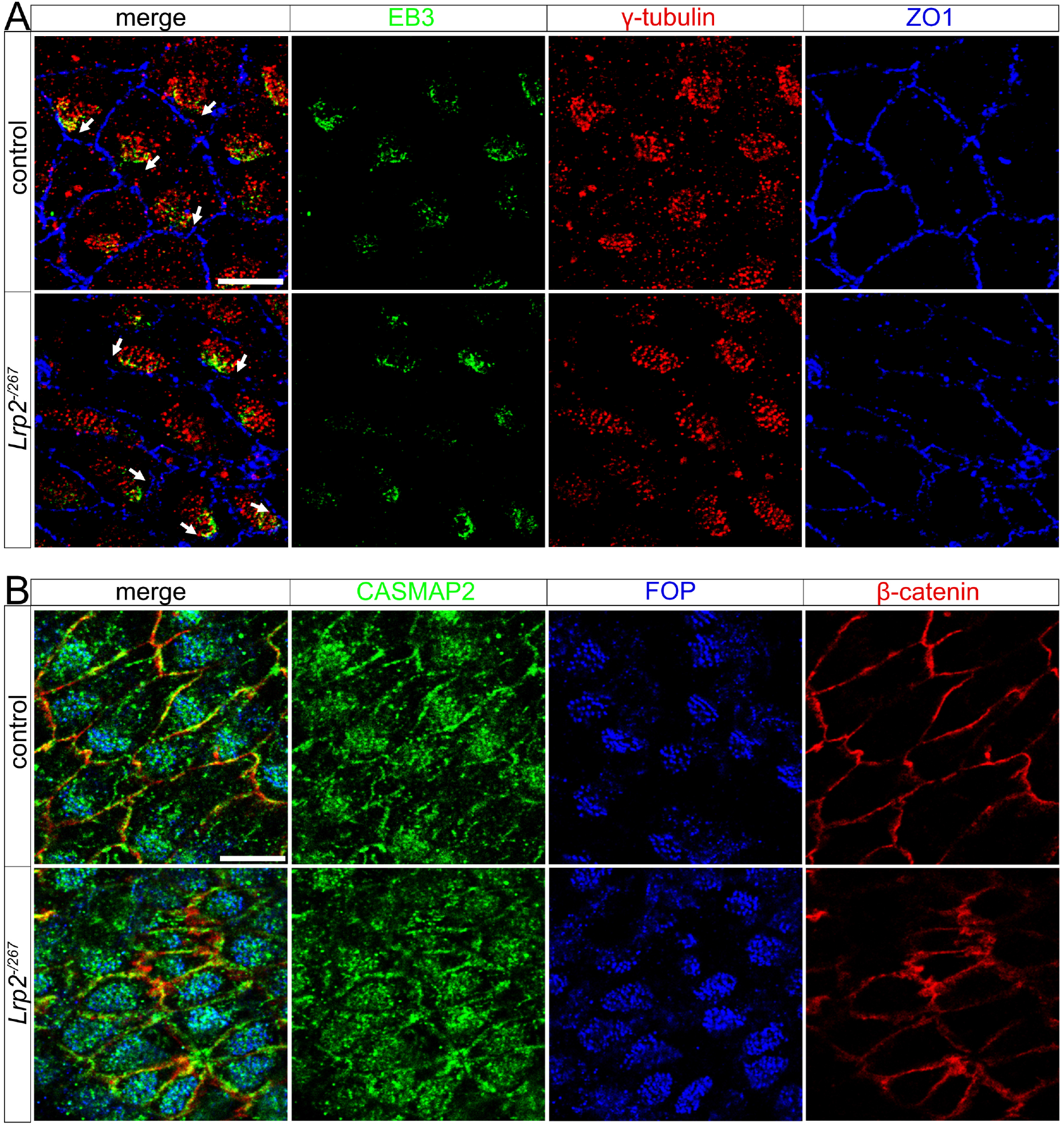
Microtubule-like structures connecting basal foot processes with the cell cortex are normal in juvenile LRP2-deficient ependyma. Immunodetection of *en face* lateral wall preparations documenting EB3 (A; green) and CASMAP2 (B; green) co-stained with γ-tubulin (red) and ZO1 (blue) or FOP (blue) and b-catenin (red), respectively. (**A**) EB3 marker of the microtubule plus end demonstrates comparable localization between control and *Lrp2^-/267^* mice, although coordinated protein localization throughout the tissue is altered (direction of white arrows) reflecting changes in translational polarity described before. (**B**) The microtubule minus end marker CASMAP2 depicts comparable protein localization between the two genotypes, but coordinated displacement of ciliary patch is impaired in the mutants as described before. Scale bar: 10 μm.

We also queried the asymmetric localization of several PCP proteins and associated factors in the ependyma, essential for the establishment of rotational polarity. Daple is a Dishevelled (Dvl)-associated protein localizing to the anterior side of ependymal cells. It is responsible for the correct positioning of the BB via cytoplasmic dynein to ensure coordinated motile cilia beating (Takagishi et al. 2020). In control ependyma, Daple localized to the anterior side of the cells, the direction to which the ciliary patch is displaced and LRP2 accumulates (Fig. 6A). Some co-localization of Daple and LRP2 close to the cell membrane at the anterior side was detected (Fig. 6A inset, arrowheads). By contrast, in ependymal cells of *Lrp2^-/267^* juvenile mice, levels of Daple were severely diminished and the residual protein showed a punctate pattern rather than the expected enrichment at the anterior cell side (Fig. 6A). Massive reduction in Daple levels as well as a failure in anterior protein accumulation was also documented in the adult *Lrp2^-/267^*ependyma (Fig. S5). Dvl1 is an intracellular adaptor protein that acts in the canonical but also in the noncanonical Wnt/PCP pathway in epithelial polarity (Wang et al. 2006; Gao 2012; Wynshaw-Boris 2012). In the ependyma of control mice, Dvl1 localized to the anterior cell side (Fig. 6B). This coordinated localization towards the ciliary patch was not seen in the *Lrp2^-/267^* ependyma (Fig. 6B). Vangl2 and Celsr1 are two proteins also implicated in directing ciliary orientation and alignment (Guirao et al. 2010; Boutin et al. 2014; Goffinet and Tissir 2017). Vangl2 and Celsr1 localize to the posterior cell side, on the opposite side of ciliary patch displacement. In line with disturbances in PCP, coordinated localization of Vangl2 and Celsr1to the posterior cell side was lost in the *Lrp2^-/267^* ependyma when compared to control tissues (Fig. 6C-D).

**Figure 6.**
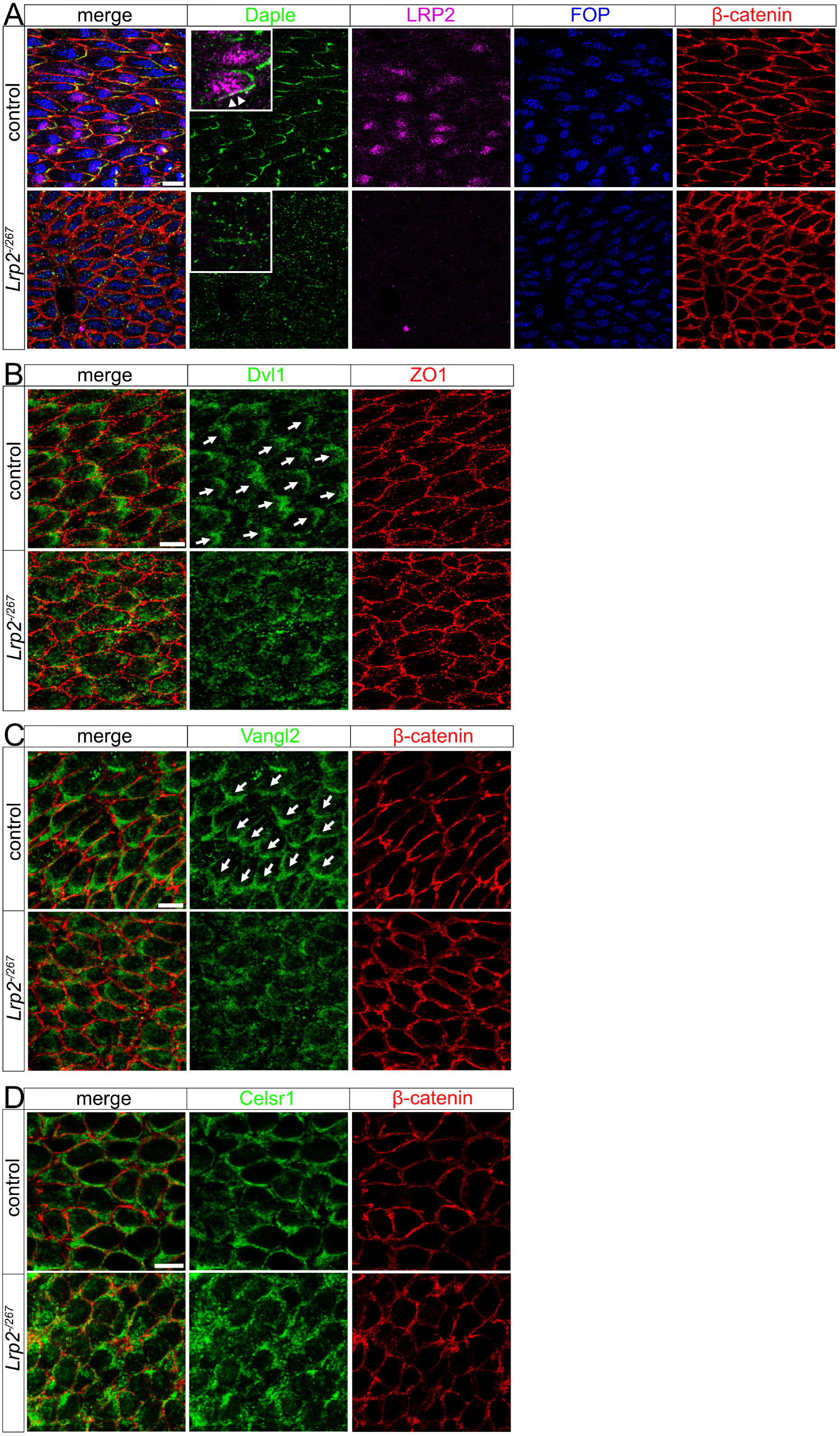
Mislocalization of planar cell polarity proteins in LRP2-deficient juvenile ependyma. **(A)** Immunohistological analyses of *en face* lateral wall preparations of the murine juvenile ependyma visualizing Daple (green), LRP2 (purple), FOP (blue), and b-catenin (red). Images are given as single or merged channel configurations. In control cells, immunosignals for Daple show an asymmetric distribution to the area of the cell to which the ciliary patch (marked by FOP) is displaced. The higher magnification inset documents close spatial proximity of LRP2 and Daple at the anterior cell surface (arrowheads). In *Lrp2^-/267^* mice, immunosignals for Daple are strongly reduced and fail to show an asymmetric distribution towards the ciliary patch. Scale bar: 10 μm. (**B-D**) Wholemount lateral wall preparations immunostained for PCP markers Dvl1 (B), Vangl2 (D), or Celsr1 (E). Cells were co-stained with apical cell surface markers ZO-1 (red) or b-catenin (red). Single channels as well as the merged channel configurations are depicted. (**B**) Dvl1 shows coordinated localization to the anterior side of the cell throughout the tissue in control ependyma (indicated by white arrows), while being unevenly distributed along the cell contour in mutants. (**C**) Vangl2 is enriched on the posterior side of the cell in a zigzag-like pattern (indicated by white arrows), a pattern lost in mutants. (**D**) similarly, Celsr1 immunosignals are enriched on the posterior side along the cell contour in control mice this clear membrane staining is lost in *Lrp2*^-/267^ mice. Scale bars: 10 μm.

One possibility explaining how LRP2 may be functionally linked to the PCP pathway components is through cytosolic adaptors that interact both with this receptor and core PCP proteins (Kowalczyk et al. 2021). This hypothesis was tested by studying the subcellular localization of the LRP2 adaptor protein Na+/H+ Exchanger Regulatory Factor 1 (NHERF1). NHERF1 binds to the cytoplasmic domain of LRP2 (Gotthardt et al. 2000) and loss of NHERF1 results in disorganized ependymal cilia (Treat et al. 2016). In the ependyma of control mice we detected NHERF1 protein in the subapical anterior cell region (Fig. 7A). Localization of NHERF1 to the side of ciliary patch displacement was lost in the ependyma of *Lrp2^-/267^* mice, documenting disruption of tissue-wide coordinated NHERF1 localization in the absence of LRP2 (Fig. 7A).

**Figure 7.**
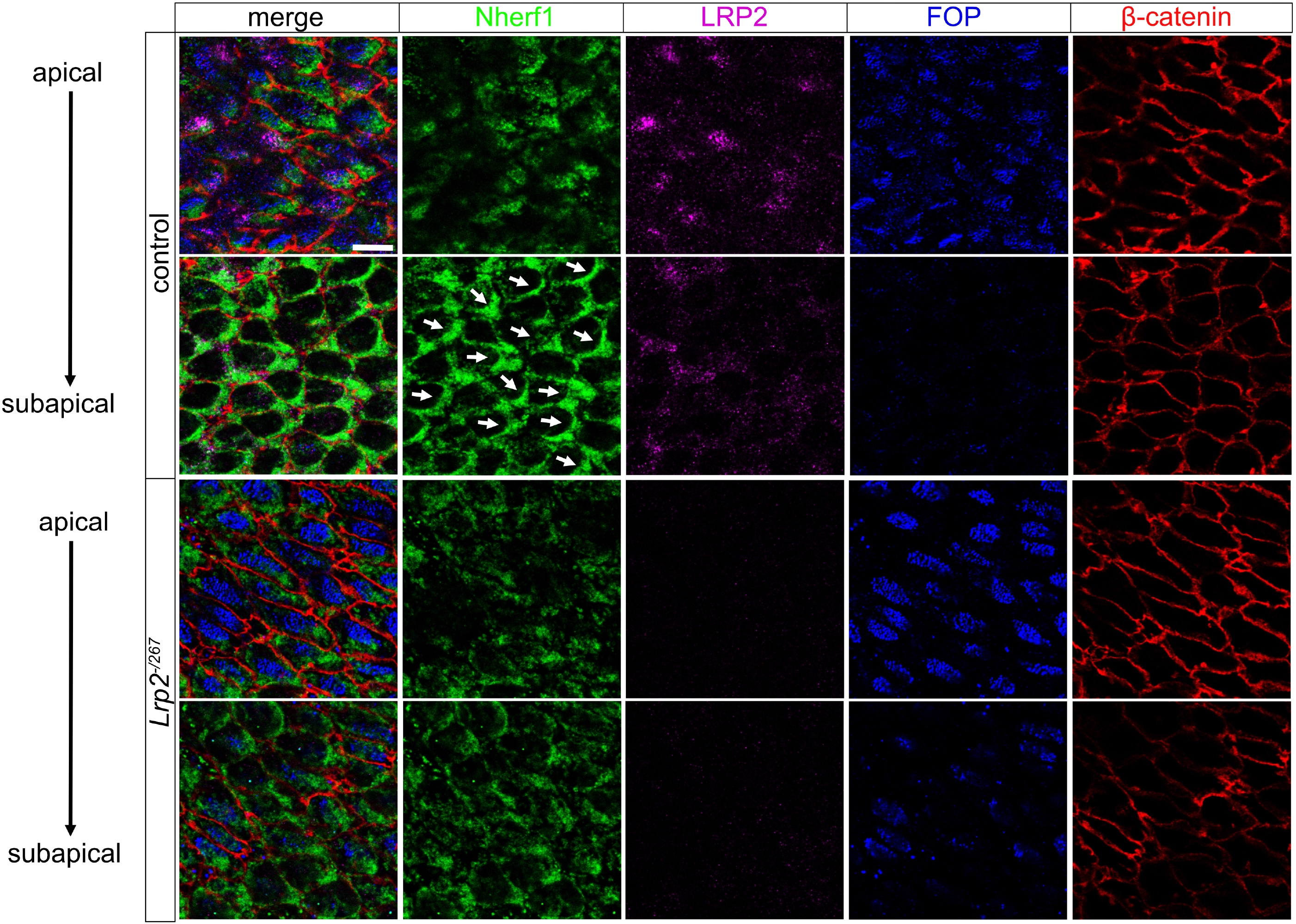
Reduction in adaptor protein NHERF1 in juvenile ependyma of LRP2-deficient mice. *En face* lateral wall preparations for NHERF1 (green), LRP2 (purple), FOP (blue), and b-catenin (red). Single projections in single channels and merged channels configurations are shown of the apical and subapical ependymal cell compartment. In control ependyma, the apical cell compartment shows colocalization of LRP2 and FOP while the subapical compartment documents asymmetric distribution of NHERF1 towards the ciliary patch, the anterior cell side (indicated by arrows). In the *Lrp2^-/267^* ependyma, localization of FOP at the apical cell compartment is normal but the polarized enrichment of NHERF1 towards the ciliary patch in the subapical compartment is lost. Scale bar: 10 μm.

## DISCUSSION

We identified LRP2 as a novel player in establishing translational and rotational polarity in the ependyma and loss of coordinated ciliary arrangement and beating in this tissue in receptor mutant mice. Polarity defects coincide with abnormal arrangement of the PCP pathway components, suggesting a role for this endocytic receptor in the organizational integrity of the PCP, possibly through its associated adaptor NHERF1.

The earliest PCP defect observed in the LRP2-deficient mice concerns failure to establish translational polarity in RGs, as evidenced by uncoordinated displacement of primary cilia (Fig. S1). This phenotype resembles defects described for loss of core PCP proteins Celsr1, Fzd3, and Vangl2 (Boutin et al. 2014). In addition, polarity of mature ependymal cells, is also affected as documented by uncoordinated displacement and absence of beating synchrony of motile cilia patches in LRP2-deficient juvenile and adult mice (Figs. 2 - 3, and S2 - S3). Such phenotypes are also shared by mice with loss of PCP pathway components, including *Vangl2* and *Fzd3* (Guirao et al. 2010; Boutin et al. 2014; Ohata et al. 2014). Impaired ciliogenesis, as documented for *Celsr2* and *Celsr2/-3* double mutant mice (Tissir et al. 2010), is likely not the reason for the polarity defects, as *Lrp2^-/267^* mice show comparable numbers of structurally and functionally intact motile cilia as control animals (Figs.1, 4, and S4).

Formally, we cannot exclude problems in induction of the passive CSF flow in the choroid plexus as the initial trigger for polarity defects in mutant RGs. However, our findings are more consistent with a role for LRP2 in sorting of PCP proteins along planar polarized microtubule structures, a process intimately linked to establishing polarity (Vladar et al. 2012; Boutin et al. 2014; Takagishi et al. 2020). Based on disrupted rotational polarity observed in *Vangl2*, *Celsr1*, and *Dvls* mouse mutants (Guirao et al. 2010; Boutin et al. 2014; Ohata et al. 2014; Yang and Mlodzik 2015), failure in proper asymmetric localization of these PCP proteins in LRP2-deficient mice (Fig. 6) is the likely cause for the uncoordinated alignment of BBs in these animals (Figs. 3 and S3). Previously, Daple, a Dvl-associated adaptor protein for cytoplasmic dynein (Redwine et al. 2017) has been shown to be involved in BB positioning by anchoring dynein to the anterior side of the cell cortex, where Dvl1 is localized. Dynein generates a pulling force on the depolymerizing microtubule connected to the basal foot, positioning the basal foot towards the anterior cell cortex and, thereby, establishing translational and rotational polarity of ependymal cells (Ishida-Takagishi et al. 2012; Takagishi et al. 2017; Takagishi et al. 2020). Daple protein levels as well as localization are severely disrupted in LRP2-deficient mice (Fig. 6 and S5), in line with the defects in ciliary patch positioning and orientation in this model.

LRP2 is mainly known as a high-capacity endocytic receptor in polarized epithelia, including the ependyma (Gajera et al. 2010). Intriguingly, loss of LRP2 activity not only impairs cellular clearance of receptor ligands but also results in breakdown of endocytic structures as shown in the renal proximal tubules of mouse mutants (Leheste et al. 1999) and Donnai-Barrow syndrome patients (Dachy et al. 2015). Based on these observations, a more general role for LRP2 as a scaffold for components of the apical vesicular trafficking machinery has been proposed (Long et al. 2022). In ependymal cells, LRP2 localizes to the ciliary pocket of motile cilia, a highly endocytic cell compartment (Molla-Herman et al. 2010; Benmerah 2013). In analogy to the kidney, breakdown of the endocytic machinery in this cell compartment seems a plausible explanation for faulty localization of PCP proteins in ependymal cells lacking LRP2. The importance of endocytic processes for PCP is exemplified by Dvl2 that interacts with the clathrin adaptor AP-2 to mediate Fzd4 internalization, an interaction required for planar cell polarity signaling (Yu et al. 2007). Also, the export of Vangl2 from the *trans* Golgi network is regulated by the clathrin adaptor AP-1 in mice (Guo et al. 2013), while in *Drosophila,* AP-1 is implicated in trafficking of Frizzled and the *Drosophila* Vangl2 orthologue Vang (Carvajal-Gonzalez et al. 2015). In line with a role of LRP2 in sorting of PCP components and associated factors along the endocytic path, LRP2 deficiency impairs apical localization of Vangl2 in the neural tube of mouse and *Xenopus* (Kowalczyk et al. 2021), as well as in the adult murine ependyma (Fig. 6).

To fulfill its role in endocytic processes, LRP2 interacts with intracellular adaptors. In the developing neural tube, recent studies implicated the cytosolic adaptor GIPC1 in LRP2-mediated sorting of Vangl2 (Kowalczyk et al. 2021). We now identified the LRP2 adaptor NHERF1 as a possible link between LRP2-mediated endocytosis and sorting of PCP proteins in the adult ependyma. NHERF1 is a cytosolic PSD-95/Drosophila disc large/Zo-1 (PDZ) adaptor localizing to the apical cell membrane in epithelia (Stemmer-Rachamimov et al. 2001), where it directly interacts with the cytoskeleton (Weinman et al. 1998). It binds to the intracellular PDZ domain of LRP2 (Gotthardt et al. 2000; Slattery et al. 2011) and its loss results in disorganized ependymal cilia, due to a failure to translocate Vangl2 to the plasma membrane (Treat et al. 2016). We now identified asymmetric localization of NHERF1 in the wild-type ependyma by sorting to the anterior side of the cell in close proximity to the ciliary patch. This asymmetric localization is lost and overall NHERF1 protein levels are reduced in the ependyma of *Lrp2^-/267^* mice (Fig. 7), suggesting a molecular mechanism whereby LRP2 links endocytosis at the apical cell compartment with sorting of PCP components.

## Acknowledgements

We are indebted to Kristin Kampf, Christine Kruse, and Maria Kahlow for expert technical assistance. The authors thank Dr. Anje Sporbert, Dr. Sandra Raimundo and Matthias Richter from the Advanced Light Microscopy Technology Platform at Max-Delbrück-Center for Molecular Medicine in the Helmholtz Association (MDC), Berlin, Germany (https://www.mdc-berlin.de/advanced-light-microscopy) for technical support and assistance with confocal microscopy.

## Author contribution

L. B., A. C., A. M., S. B., E. I. C. performed experiments and evaluated data. A. C., C. B., T. E. W. conceived experiments and evaluated data. M. M. provided reagents. A. C. and T. E. W. wrote the manuscript.

## Funding

Studies were funded in part by a grant from the Deutsche Forschungsgemeinschaft (CH 1838/1-1) to AC

## DECLARATIONS

### Conflict of Interests

The authors declare no competing interests.

### Ethical Approval

All experiments involving animals were performed according to institutional guidelines following approval by local authorities (G0042/19).

### Informed Consent

Not Applicable

## Notes

### Competing Interest Statement

The authors have declared no competing interest.

## REFERENCES

Akhmanova A, Hoogenraad CC (2015) Microtubule minus-end-targeting proteins. Curr Biol 25:R162–171. https://doi.org/10.1016/j.cub.2014.12.027

Argraves WS, Morales CR (2004) Immunolocalization of cubilin, megalin, apolipoprotein J, and apolipoprotein A-I in the uterus and oviduct. Mol Reprod Dev 69:419–427. https://doi.org/10.1002/mrd.20174

Benmerah A (2013) The ciliary pocket. Current Opinion in Cell Biology 25:78–84. https://doi.org/10.1016/j.ceb.2012.10.011

Boutin C, Labedan P, Dimidschstein J, Richard F, Cremer H, André P, Yang Y, Montcouquiol M, Goffinet AM, Tissir F (2014) A dual role for planar cell polarity genes in ciliated cells. Proceedings of the National Academy of Sciences 111:E3129– E3138. https://doi.org/10.1073/pnas.1404988111

Butler MT, Wallingford JB (2017) Planar cell polarity in development and disease. Nat Rev Mol Cell Biol 18:375–388. https://doi.org/10.1038/nrm.2017.11

Carvajal-Gonzalez JM, Balmer S, Mendoza M, Dussert A, Collu G, Roman A-C, Weber U, Ciruna B, Mlodzik M (2015) The clathrin adaptor AP-1 complex and Arf1 regulate planar cell polarity in vivo. Nat Commun 6:6751. https://doi.org/10.1038/ncomms7751

Christ A, Christa A, Kur E, Lioubinski O, Bachmann S, Willnow TE, Hammes A (2012) LRP2 is an auxiliary SHH receptor required to condition the forebrain ventral midline for inductive signals. Dev Cell 22:268–278. https://doi.org/10.1016/j.devcel.2011.11.023

Christ A, Herzog K, Willnow TE (2016) LRP2, an auxiliary receptor that controls sonic hedgehog signaling in development and disease. Dev Dyn 245:569–579. https://doi.org/10.1002/dvdy.24394

Christensen EI, Nielsen S, Moestrup SK, Borre C, Maunsbach AB, de Heer E, Ronco P, Hammond TG, Verroust P (1995) Segmental distribution of the endocytosis receptor gp330 in renal proximal tubules. Eur J Cell Biol 66:349–364

Dachy A, Paquot F, Debray G, Bovy C, Christensen EI, Collard L, Jouret F (2015) In-depth phenotyping of a Donnai–Barrow patient helps clarify proximal tubule dysfunction. Pediatr Nephrol 30:1027–1031. https://doi.org/10.1007/s00467-014-3037-7

Gajera CR, Emich H, Lioubinski O, Christ A, Beckervordersandforth-Bonk R, Yoshikawa K, Bachmann S, Christensen EI, Götz M, Kempermann G, Peterson AS, Willnow TE, Hammes A (2010) LRP2 in ependymal cells regulates BMP signaling in the adult neurogenic niche. J Cell Sci 123:1922–1930. https://doi.org/10.1242/jcs.065912

Gao B (2012) Wnt regulation of planar cell polarity (PCP). Curr Top Dev Biol 101:263–295. https://doi.org/10.1016/B978-0-12-394592-1.00008-9

Geraldo S, Khanzada UK, Parsons M, Chilton JK, Gordon-Weeks PR (2008) Targeting of the F-actin-binding protein drebrin by the microtubule plus-tip protein EB3 is required for neuritogenesis. Nat Cell Biol 10:1181–1189. https://doi.org/10.1038/ncb1778

Goffinet AM, Tissir F (2017) Seven pass Cadherins CELSR1-3. Semin Cell Dev Biol 69:102–110. https://doi.org/10.1016/j.semcdb.2017.07.014

Gotthardt M, Trommsdorff M, Nevitt MF, Shelton J, Richardson JA, Stockinger W, Nimpf J, Herz J (2000) Interactions of the low density lipoprotein receptor gene family with cytosolic adaptor and scaffold proteins suggest diverse biological functions in cellular communication and signal transduction. J Biol Chem 275:25616–25624. https://doi.org/10.1074/jbc.M000955200

Guirao B, Meunier A, Mortaud S, Aguilar A, Corsi J-M, Strehl L, Hirota Y, Desoeuvre A, Boutin C, Han Y-G, Mirzadeh Z, Cremer H, Montcouquiol M, Sawamoto K, Spassky N (2010) Coupling between hydrodynamic forces and planar cell polarity orients mammalian motile cilia. Nat Cell Biol 12:341–350. https://doi.org/10.1038/ncb2040

Guo Y, Zanetti G, Schekman R (2013) A novel GTP-binding protein-adaptor protein complex responsible for export of Vangl2 from the trans Golgi network. Elife 2:e00160. https://doi.org/10.7554/eLife.00160

Hagiwara H, Kano A, Aoki T, Ohwada N, Takata K (2000) Localization of gamma-tubulin to the basal foot associated with the basal body extending a cilium. Histochem J 32:669–671. https://doi.org/10.1023/a:1004163315822

Harrison C, Shao H, Strutt H, Strutt D (2020) Molecular mechanisms mediating asymmetric subcellular localisation of the core planar polarity pathway proteins. Biochem Soc Trans 48:1297–1308. https://doi.org/10.1042/BST20190404

Hirota Y, Meunier A, Huang S, Shimozawa T, Yamada O, Kida YS, Inoue M, Ito T, Kato H, Sakaguchi M, Sunabori T, Nakaya M-A, Nonaka S, Ogura T, Higuchi H, Okano H, Spassky N, Sawamoto K (2010) Planar polarity of multiciliated ependymal cells involves the anterior migration of basal bodies regulated by non-muscle myosin II. Development 137:3037–3046. https://doi.org/10.1242/dev.050120

Hyland RM, Brody SL (2021) Impact of Motile Ciliopathies on Human Development and Clinical Consequences in the Newborn. Cells 11:125. https://doi.org/10.3390/cells11010125

Ishida-Takagishi M, Enomoto A, Asai N, Ushida K, Watanabe T, Hashimoto T, Kato T, Weng L, Matsumoto S, Asai M, Murakumo Y, Kaibuchi K, Kikuchi A, Takahashi M (2012) The Dishevelled-associating protein Daple controls the non-canonical Wnt/Rac pathway and cell motility. Nat Commun 3:859. https://doi.org/10.1038/ncomms1861

Jaworski J, Kapitein LC, Gouveia SM, Dortland BR, Wulf PS, Grigoriev I, Camera P, Spangler SA, Di Stefano P, Demmers J, Krugers H, Defilippi P, Akhmanova A, Hoogenraad CC (2009) Dynamic microtubules regulate dendritic spine morphology and synaptic plasticity. Neuron 61:85–100. https://doi.org/10.1016/j.neuron.2008.11.013

Kantarci S, Al-Gazali L, Hill RS, Donnai D, Black GCM, Bieth E, Chassaing N, Lacombe D, Devriendt K, Teebi A, Loscertales M, Robson C, Liu T, MacLaughlin DT, Noonan KM, Russell MK, Walsh CA, Donahoe PK, Pober BR (2007) Mutations in LRP2, which encodes the multiligand receptor megalin, cause Donnai-Barrow and facio-oculo-acoustico-renal syndromes. Nat Genet 39:957–959. https://doi.org/10.1038/ng2063

Kapania EM, Stern BM, Sharma G (2022) Ciliary Dysfunction. In: StatPearls. StatPearls Publishing, Treasure Island (FL)

Kowalczyk I, Lee C, Schuster E, Hoeren J, Trivigno V, Riedel L, Görne J, Wallingford JB, Hammes A, Feistel K (2021) Neural tube closure requires the endocytic receptor Lrp2 and its functional interaction with intracellular scaffolds. Development 148:dev195008. https://doi.org/10.1242/dev.195008

Kumar V, Umair Z, Kumar S, Goutam RS, Park S, Kim J (2021) The regulatory roles of motile cilia in CSF circulation and hydrocephalus. Fluids Barriers CNS 18:31. https://doi.org/10.1186/s12987-021-00265-0

Kunimoto K, Yamazaki Y, Nishida T, Shinohara K, Ishikawa H, Hasegawa T, Okanoue T, Hamada H, Noda T, Tamura A, Tsukita S, Tsukita S (2012) Coordinated ciliary beating requires Odf2-mediated polarization of basal bodies via basal feet. Cell 148:189–200. https://doi.org/10.1016/j.cell.2011.10.052

Kur E, Mecklenburg N, Cabrera RM, Willnow TE, Hammes A (2014) LRP2 mediates folate uptake in the developing neural tube. J Cell Sci 127:2261–2268. https://doi.org/10.1242/jcs.140145

Labedan P, Matthews C, Kodjabachian L, Cremer H, Tissir F, Boutin C (2016) Dissection and Staining of Mouse Brain Ventricular Wall for the Analysis of Ependymal Cell Cilia Organization. Bio-protocol 6:e1757–e1757

Leheste JR, Rolinski B, Vorum H, Hilpert J, Nykjaer A, Jacobsen C, Aucouturier P, Moskaug JO, Otto A, Christensen EI, Willnow TE (1999) Megalin knockout mice as an animal model of low molecular weight proteinuria. Am J Pathol 155:1361–1370

Liu H, Zheng J, Zhu L, Xie L, Chen Y, Zhang Y, Zhang W, Yin Y, Peng C, Zhou J, Zhu X, Yan X (2021) Wdr47, Camsaps, and Katanin cooperate to generate ciliary central microtubules. Nat Commun 12:5796. https://doi.org/10.1038/s41467-021-26058-5

Long KR, Rbaibi Y, Bondi CD, Ford BR, Poholek AC, Boyd-Shiwarski CR, Tan RJ, Locker JD, Weisz OA (2022) Cubilin-, megalin-, and Dab2-dependent transcription revealed by CRISPR/Cas9 knockout in kidney proximal tubule cells. Am J Physiol Renal Physiol 322:F14–F26. https://doi.org/10.1152/ajprenal.00259.2021

Lundgren S, Carling T, Hjälm G, Juhlin C, Rastad J, Pihlgren U, Rask L, Akerström G, Hellman P (1997) Tissue distribution of human gp330/megalin, a putative Ca(2+)- sensing protein. J Histochem Cytochem 45:383–392

Marshall WF, Kintner C (2008) Cilia orientation and the fluid mechanics of development. Curr Opin Cell Biol 20:48–52. https://doi.org/10.1016/j.ceb.2007.11.009

Mirzadeh Z, Han Y-G, Soriano-Navarro M, García-Verdugo JM, Alvarez-Buylla A (2010) Cilia organize ependymal planar polarity. J Neurosci 30:2600–2610. https://doi.org/10.1523/JNEUROSCI.3744-09.2010

Mirzadeh Z, Merkle FT, Soriano-Navarro M, Garcia-Verdugo JM, Alvarez-Buylla A (2008) Neural stem cells confer unique pinwheel architecture to the ventricular surface in neurogenic regions of the adult brain. Cell Stem Cell 3:265–278. https://doi.org/10.1016/j.stem.2008.07.004

Miura K (2020) Bleach correction ImageJ plugin for compensating the photobleaching of time-lapse sequences. F1000Res 9:1494. https://doi.org/10.12688/f1000research.27171.1

Molla-Herman A, Ghossoub R, Blisnick T, Meunier A, Serres C, Silbermann F, Emmerson C, Romeo K, Bourdoncle P, Schmitt A, Saunier S, Spassky N, Bastin P, Benmerah A (2010) The ciliary pocket: an endocytic membrane domain at the base of primary and motile cilia. J Cell Sci 123:1785–1795. https://doi.org/10.1242/jcs.059519

Nykjaer A, Willnow TE (2002) The low-density lipoprotein receptor gene family: a cellular Swiss army knife? Trends Cell Biol 12:273–280

Ohata S, Alvarez-Buylla A (2016) Planar Organization of Multiciliated Ependymal (E1) Cells in the Brain Ventricular Epithelium. Trends in Neurosciences 39:543–551. https://doi.org/10.1016/j.tins.2016.05.004

Ohata S, Nakatani J, Herranz-Pérez V, Cheng J, Belinson H, Inubushi T, Snider WD, García-Verdugo JM, Wynshaw-Boris A, Alvarez-Buylla A (2014) Loss of Dishevelleds disrupts planar polarity in ependymal motile cilia and results in hydrocephalus. Neuron 83:558–571. https://doi.org/10.1016/j.neuron.2014.06.022

Park TJ, Mitchell BJ, Abitua PB, Kintner C, Wallingford JB (2008) Dishevelled controls apical docking and planar polarization of basal bodies in ciliated epithelial cells. Nat Genet 40:871–879. https://doi.org/10.1038/ng.104

Redmond SA, Figueres-Oñate M, Obernier K, Nascimento MA, Parraguez JI, López-Mascaraque L, Fuentealba LC, Alvarez-Buylla A (2019) Development of Ependymal and Postnatal Neural Stem Cells and Their Origin from a Common Embryonic Progenitor. Cell Rep 27:429–441.e3. https://doi.org/10.1016/j.celrep.2019.01.088

Redwine WB, DeSantis ME, Hollyer I, Htet ZM, Tran PT, Swanson SK, Florens L, Washburn MP, Reck-Peterson SL (2017) The human cytoplasmic dynein interactome reveals novel activators of motility. Elife 6:e28257. https://doi.org/10.7554/eLife.28257

Robinson AM, Takahashi S, Brotslaw EJ, Ahmad A, Ferrer E, Procissi D, Richter C-P, Cheatham MA, Mitchell BJ, Zheng J (2020) CAMSAP3 facilitates basal body polarity and the formation of the central pair of microtubules in motile cilia. Proc Natl Acad Sci U S A 117:13571–13579. https://doi.org/10.1073/pnas.1907335117

Rosenfeld JA, Ballif BC, Martin DM, Aylsworth AS, Bejjani BA, Torchia BS, Shaffer LG (2010) Clinical characterization of individuals with deletions of genes in holoprosencephaly pathways by aCGH refines the phenotypic spectrum of HPE. Hum Genet. https://doi.org/10.1007/s00439-009-0778-7

Sabatino JA, Stokes BA, Zohn IE (2017) Prevention of neural tube defects in Lrp2 mutant mouse embryos by folic acid supplementation. Birth Defects Res 109:16–26. https://doi.org/10.1002/bdra.23589

Satir P, Heuser T, Sale WS (2014) A Structural Basis for How Motile Cilia Beat. Bioscience 64:1073–1083. https://doi.org/10.1093/biosci/biu180

Slattery C, Jenkin KA, Lee A, Simcocks AC, McAinch AJ, Poronnik P, Hryciw DH (2011) Na+-H+ exchanger regulatory factor 1 (NHERF1) PDZ scaffold binds an internal binding site in the scavenger receptor megalin. Cell Physiol Biochem 27:171–178. https://doi.org/10.1159/000325219

Spoelgen R, Hammes A, Anzenberger U, Zechner D, Andersen OM, Jerchow B, Willnow TE (2005) LRP2/megalin is required for patterning of the ventral telencephalon. Development 132:405–414. https://doi.org/10.1242/dev.01580

Stemmer-Rachamimov AO, Wiederhold T, Nielsen GP, James M, Pinney-Michalowski D, Roy JE, Cohen WA, Ramesh V, Louis DN (2001) NHE-RF, a merlin-interacting protein, is primarily expressed in luminal epithelia, proliferative endometrium, and estrogen receptor-positive breast carcinomas. Am J Pathol 158:57–62. https://doi.org/10.1016/S0002-9440(10)63944-2

Takagishi M, Esaki N, Takahashi K, Takahashi M (2020) Cytoplasmic Dynein Functions in Planar Polarization of Basal Bodies within Ciliated Cells. iScience 23:101213. https://doi.org/10.1016/j.isci.2020.101213

Takagishi M, Sawada M, Ohata S, Asai N, Enomoto A, Takahashi K, Weng L, Ushida K, Ara H, Matsui S, Kaibuchi K, Sawamoto K, Takahashi M (2017) Daple Coordinates Planar Polarized Microtubule Dynamics in Ependymal Cells and Contributes to Hydrocephalus. Cell Rep 20:960–972. https://doi.org/10.1016/j.celrep.2017.06.089

Tissir F, Qu Y, Montcouquiol M, Zhou L, Komatsu K, Shi D, Fujimori T, Labeau J, Tyteca D, Courtoy P, Poumay Y, Uemura T, Goffinet AM (2010) Lack of cadherins Celsr2 and Celsr3 impairs ependymal ciliogenesis, leading to fatal hydrocephalus. Nat Neurosci 13:700–707. https://doi.org/10.1038/nn.2555

Treat AC, Wheeler DS, Stolz DB, Tsang M, Friedman PA, Romero G (2016) The PDZ Protein Na+/H+ Exchanger Regulatory Factor-1 (NHERF1) Regulates Planar Cell Polarity and Motile Cilia Organization. PLoS One 11:e0153144. https://doi.org/10.1371/journal.pone.0153144

Vladar EK, Bayly RD, Sangoram AM, Scott MP, Axelrod JD (2012) Microtubules enable the planar cell polarity of airway cilia. Curr Biol 22:2203–2212. https://doi.org/10.1016/j.cub.2012.09.046

Wallingford JB (2010) Planar cell polarity signaling, cilia and polarized ciliary beating. Curr Opin Cell Biol 22:597–604. https://doi.org/10.1016/j.ceb.2010.07.011

Wang J, Hamblet NS, Mark S, Dickinson ME, Brinkman BC, Segil N, Fraser SE, Chen P, Wallingford JB, Wynshaw-Boris A (2006) Dishevelled genes mediate a conserved mammalian PCP pathway to regulate convergent extension during neurulation. Development 133:1767–1778. https://doi.org/10.1242/dev.02347

Weinman EJ, Steplock D, Tate K, Hall RA, Spurney RF, Shenolikar S (1998) Structure-function of recombinant Na/H exchanger regulatory factor (NHE-RF). J Clin Invest 101:2199–2206. https://doi.org/10.1172/JCI204

Werner ME, Hwang P, Huisman F, Taborek P, Yu CC, Mitchell BJ (2011) Actin and microtubules drive differential aspects of planar cell polarity in multiciliated cells. Journal of Cell Biology 195:19–26. https://doi.org/10.1083/jcb.201106110

Willnow TE, Christ A (2017) Endocytic receptor LRP2/megalin—of holoprosencephaly and renal Fanconi syndrome. Pflugers Arch - Eur J Physiol 469:907–916. https://doi.org/10.1007/s00424-017-1992-0

Willnow TE, Hilpert J, Armstrong SA, Rohlmann A, Hammer RE, Burns DK, Herz J (1996) Defective forebrain development in mice lacking gp330/megalin. Proc Natl Acad Sci USA 93:8460–8464

Worthington WC, Cathcart RS (1963) Ependymal Cilia: Distribution and Activity in the Adult Human Brain. Science 139:221–222. https://doi.org/10.1126/science.139.3551.221

Wynshaw-Boris A (2012) Dishevelled: in vivo roles of a multifunctional gene family during development. Curr Top Dev Biol 101:213–235. https://doi.org/10.1016/B978-0-12-394592-1.00007-7

Yang Y, Mlodzik M (2015) Wnt-Frizzled/planar cell polarity signaling: cellular orientation by facing the wind (Wnt). Annu Rev Cell Dev Biol 31:623–646. https://doi.org/10.1146/annurev-cellbio-100814-125315

Yu A, Rual J-F, Tamai K, Harada Y, Vidal M, He X, Kirchhausen T (2007) Association of Dishevelled with the clathrin AP-2 adaptor is required for Frizzled endocytosis and planar cell polarity signaling. Dev Cell 12:129–141. https://doi.org/10.1016/j.devcel.2006.10.015

Zarbalis K, May SR, Shen Y, Ekker M, Rubenstein JLR, Peterson AS (2004) A focused and efficient genetic screening strategy in the mouse: identification of mutations that disrupt cortical development. PLoS Biol 2:E219. https://doi.org/10.1371/journal.pbio.0020219

