## Supplementary Information for "LRP2 contributes to planar cell polarity-dependent coordination of motile cilia function"

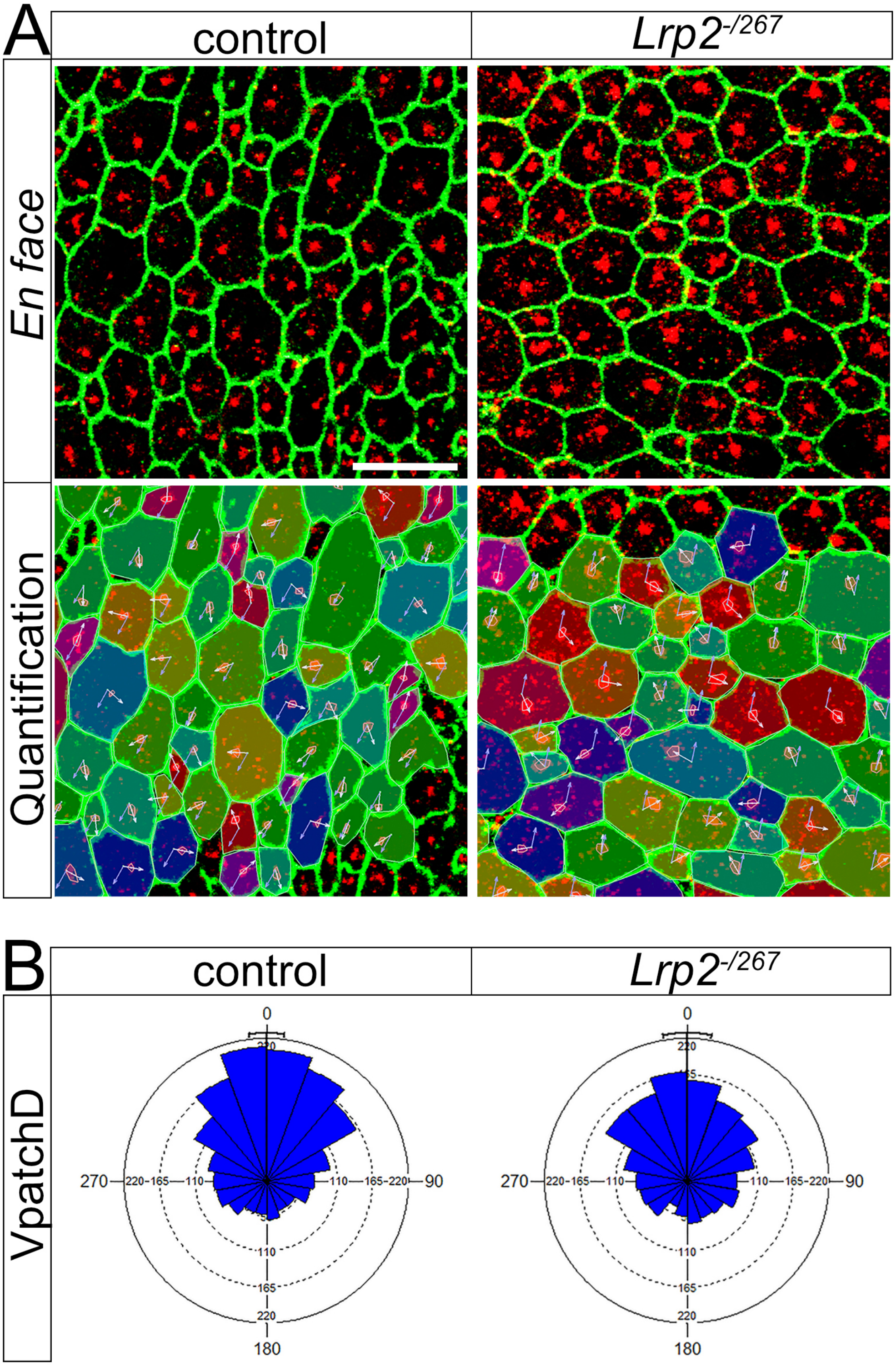


Bunatyan et al., Figure S1

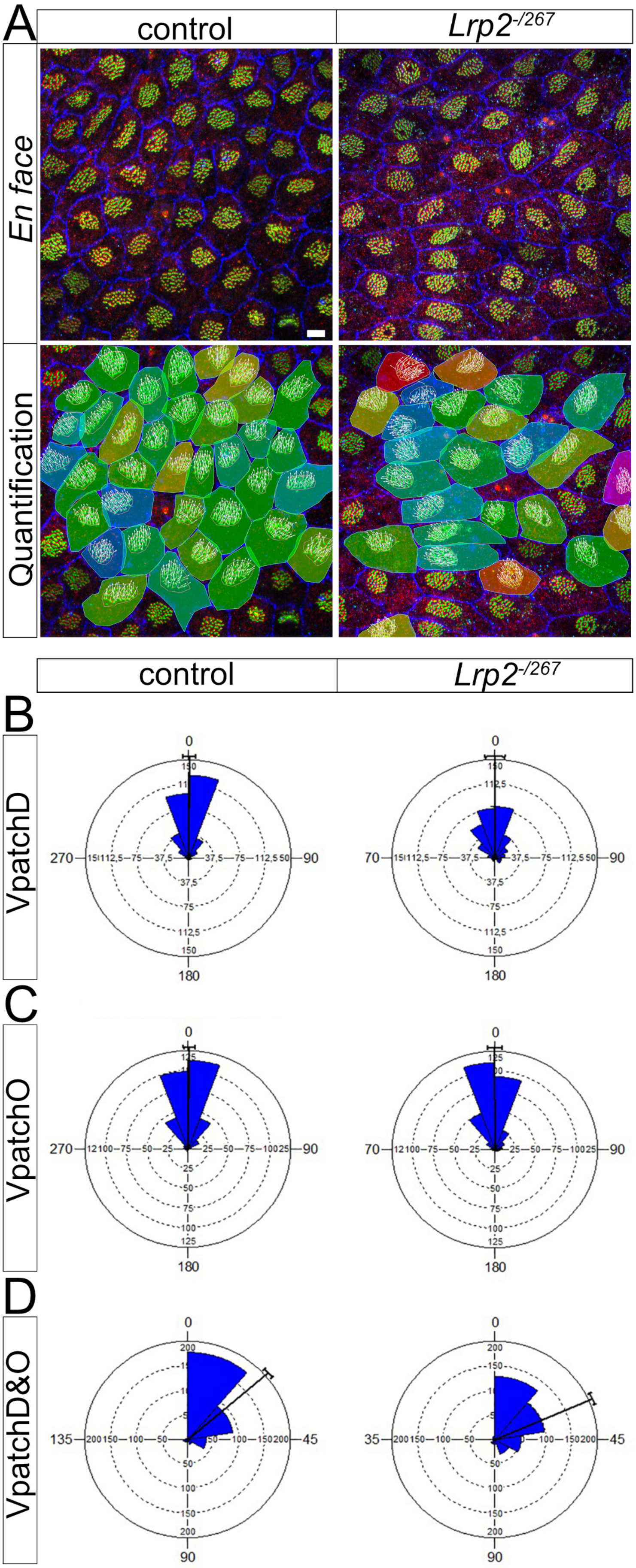


Bunatyan et al., Figure S2

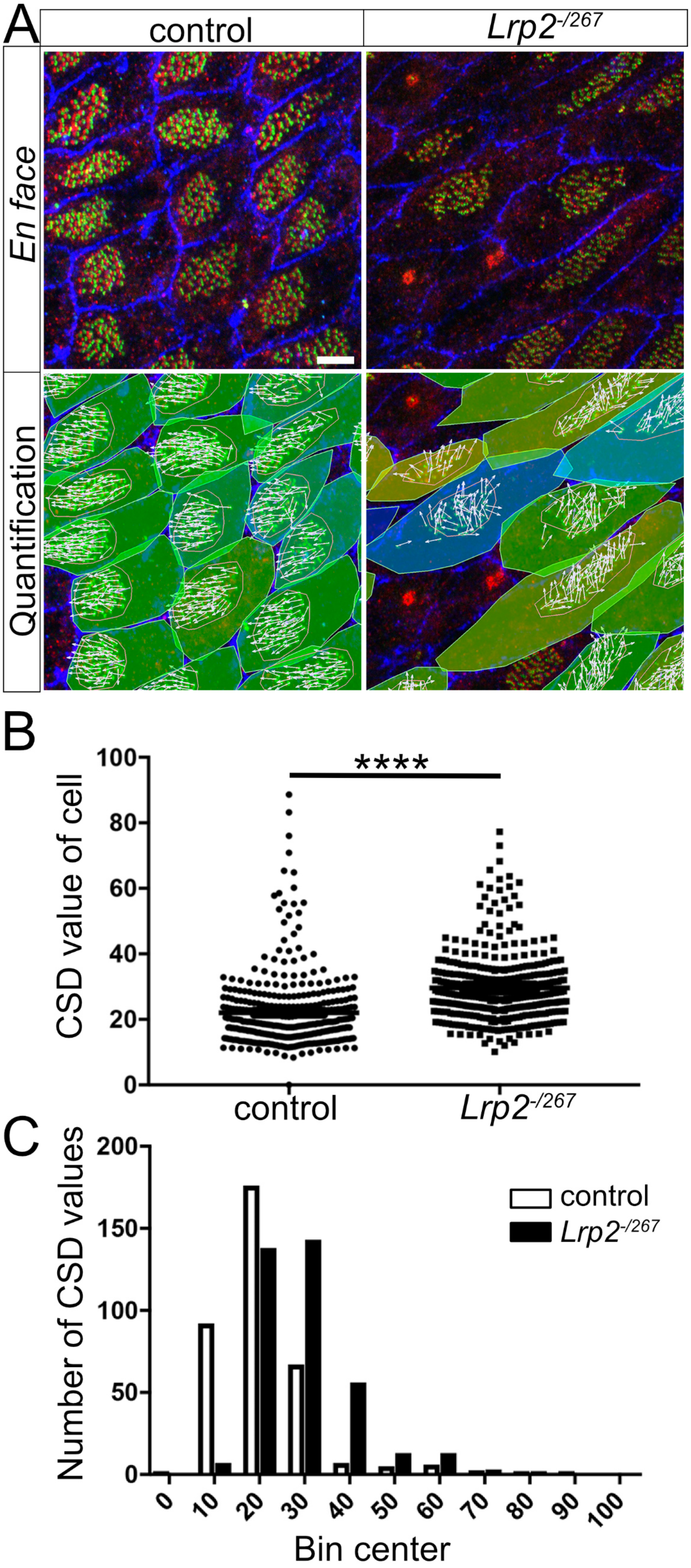


Bunatyan et al., Figure S3

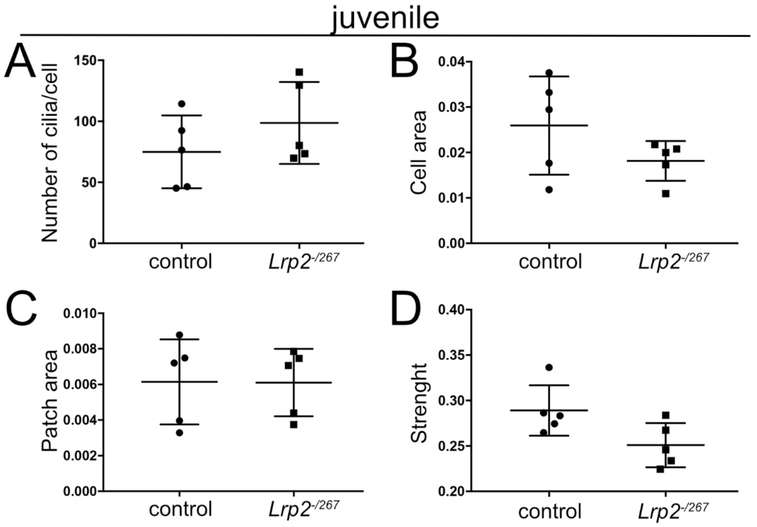


Bunatyan et al., Figure S4

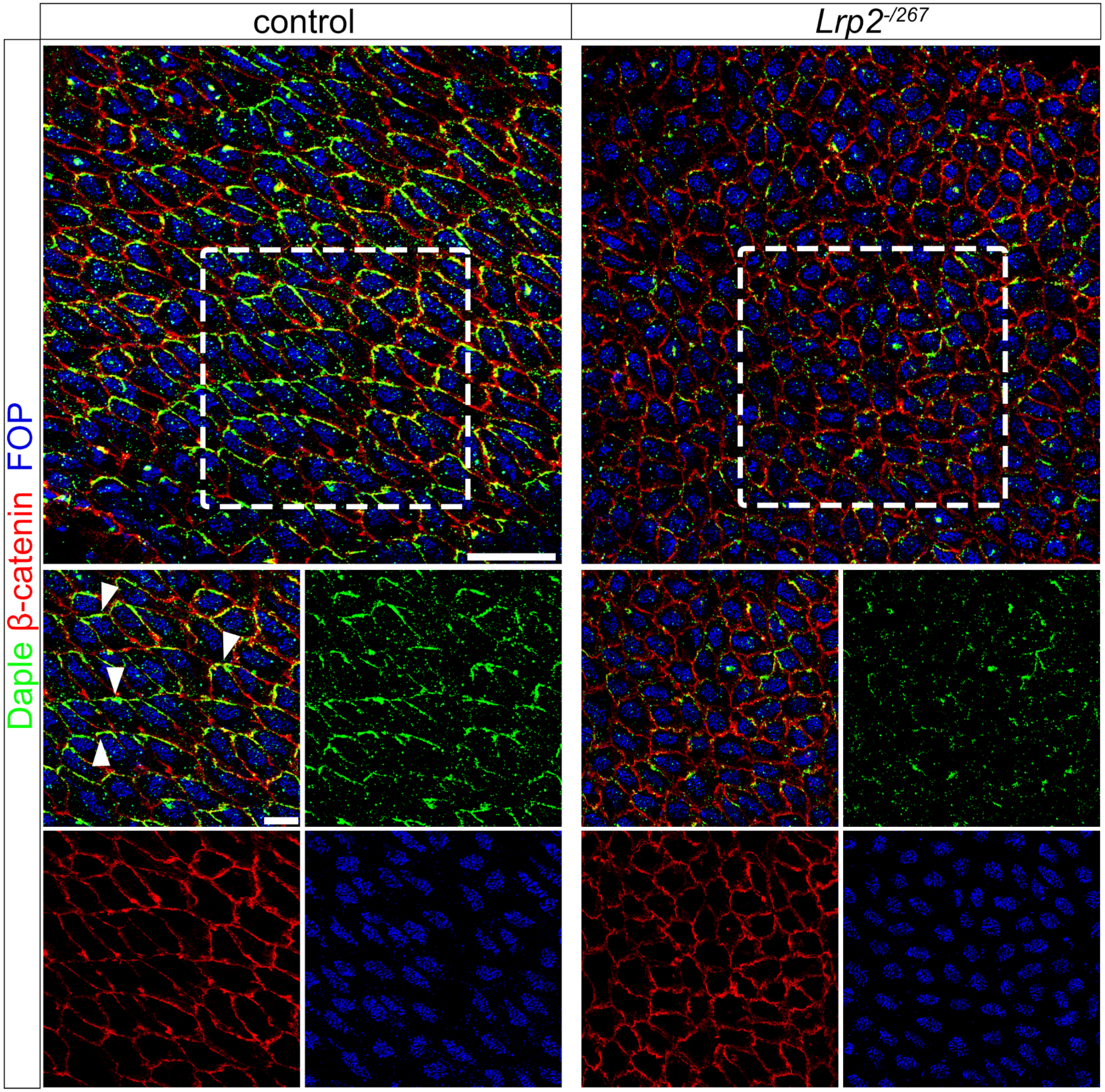


Bunatyan et al., Figure S5

### SUPPLEMENTARY FIGURE LEGENDS

**Figure S1. Disturbed translational polarity in LRP2-deficient ependymal radial progenitors**

(**A**) The upper panels depict immunodetections of basal body marker γ-tubulin (red) and ZO1 (green) marking primary cilia and apical cell boundaries on *en face* preparations of the lateral wall of newborn control and *Lrp2*$^{-/267}$ mice. Scale bar: 10 µm. The lower panels show same images with visualization of tissue-wide translational polarity based on displacement of the primary cilium from cell center using the Biotool1 software (github.com/pol51/biotool1). White arrows represent vectors of basal body displacement while blue arrows indicate the mean vectors of displacement across the tissue. Cells with misaligned primary cilium displacement compared to the mean across the tissue are marked by red to violet colour spectra. (**B**) Graphical representation of circular statistical analysis using Watson $U^2$ test (control: 1872 cells, *Lrp2*$^{-/267}$: 1741 cells, 5 animals per genotype) documenting a significantly broader distribution of VpatchD angles around the mean in *Lrp2*$^{-/267}$ ependymal cells as compared with control cells $p<0.001$).

**Figure S2. Misalignment of ciliary patch displacement and beating orientation in the LRP2-deficient adult ependyma**

**(A)** The upper panels show *en face* views of the ventricular lateral wall of adult control and *Lrp2*$^{-/267}$ mice (postnatal day >70) immuno-stained for basal body markers FOP (green) and γ-tubulin (red), as well as for apical cell surface marker ZO1 (blue). Scale bar: 5 µm. The lower panels depict the same images visualizing analysis of tissue-wide planar cell polarity using the Biotool1 software. Arrows indicate the vectors from FOP to γ-tubulin immunosignals in individual cilia, defined as Vcil. Color coding of individual cells describes the degree of ciliary patch displacement relative to the centre of the cell. The stronger the deviation of patch displacement of individual cells from the average vector of the entire field, the further the color shifts from green towards the red or blue colour spectra. (**B-D**) Graphical representation of circular statistical analysis using Watson $U^2$ test (controls: 358 cells, *Lrp2*$^{-/267}$ mutants: 374 cells, 5 animals per genotype) documenting impaired coordination of ciliary patch displacement (VpatchD, $p<0.001$; B) as well as beating orientation (VpatchO, $p<0.05$; C) in receptor-deficient as compared to control cells. Also, alignment of patch displacement and beating orientation (VpatchD&O; D) is significantly reduced in mutants as shown by Watson $U^2$ test (controls: 337 cells, *Lrp2*$^{-/267}$: 342 cells, 5 animals per genotype; $p<0.001$).

**Figure S3. Uncoordinated beating orientation of individual cilia within a ciliary patch of the LRP2-deficient adult ependyma**

(**A**) The upper panels depict *en face* preparations of the LW of adult control and *Lrp2*$^{-/267}$ mice (P>70) stained for FOP (green), γ-tubulin (red), and ZO1 (blue). Scale bar: 5 µm. The lower panels depict the same images visualizing individual cilia beating direction (direction of arrow) projected on ependymal cells. Arrows indicate the vector from FOP to γ-tubulin signals and are defined by Vcil. (**B**) Vcils are used to determine the beating coordination of cilia in single cells given by circular standard deviation (CSD) value. CSD values are significantly lower in control (391 cells from 5 different animals, one dot per cell) as compared to *Lrp2*$^{-/267}$ mice (363 cells from 5 different animals; unpaired t test, $p<0.0001$). (**C**) Random beating orientation in mutant cells is also reflected by the frequency interval demonstrating more control cells (y-axis) with lower CSD value (x-axis) as compared to *Lrp2*$^{-/267}$ cells.

**Figure S4. Ciliary patch organization in ependymal cells of LRP2-deficient juvenile mice**

Using Biotool1 software and analyzing 5 animals per genotype (265 cells in control and 284 cells in *Lrp2*$^{-/267}$ mice), no significant changes are detected in (**A**) the number of cilia per cell, in (**B**) cell surface area (Cell area), in (**C**) ciliary patch size (Patch area) and in (**D**) displacement of the ciliary patch relative to the cell center (Strenght) between the two analyzed genotypes.

**Figure S5. Reduced and mislocalized Daple protein in the adult ependyma of LRP2-deficient mice**

Whole mount immunodetection of LW preparations are stained for Daple (green), b-catenin (red) and FOP (blue). Merged overview pictures are shown for both genotypes. Scale bar: 25 µm. Detailed comparison of Daple protein immunosignals are given in single as well as merged channel configuration in higher magnification images. Polarized Daple distribution is detected in control mice (arrowheads). Significant protein reduction as well as lost coordinated protein distribution is demonstrated in *Lrp2*$^{-/267}$ mice. Scale bar: 10 µm.
